# The aging-induced long non-coding RNA *MIRIAL* controls endothelial cell and mitochondrial function

**DOI:** 10.1101/2024.02.28.582649

**Authors:** Caroline Kohnle, Susanne Koziarek, Timothy Warwick, Kosta Theodorou, Ariane Fischer, Rio Putra Juni, Marion Muhly-Reinholz, Denise Busscher, Frederike Lam, Felix Vetter, Julian U. G. Wagner, Janina Sommer, Anna Theresa Gimbel, Laura Stanicek, Stefan Günther, Ilka Wittig, Lars Maegdefessel, Matthias S. Leisegang, Ralf P. Brandes, Stefanie Dimmeler, Reinier A. Boon

## Abstract

**Aims:** Vascular aging is characterized by the progressive deterioration of endothelial function. Long non-coding RNAs (lncRNAs) are critical regulators of gene expression and protein function. However, their involvement in aging-related dysregulation of endothelial cell function remains largely unknown. Here, we aim to characterize the aging-regulated lncRNA *MIRIAL* in endothelial cells.

**Methods + Results:** We identified *Mirial* as an aging-induced lncRNA in RNA-sequencing data of mouse cardiac endothelial cells. In human umbilical vein endothelial cells (HUVECs), gapmer-mediated knockdown of *MIRIAL* led to decreases in proliferation, migration and basal angiogenic sprouting. Additionally, *MIRIAL* knockdown led to increased mitochondrial mass, spare respiratory capacity, and vascular endothelial growth factor (VEGF)-stimulated sprouting. Mechanistically, we demonstrate that *MIRIAL* forms an *RNA*·DNA:DNA triple helix (triplex) with a regulatory region of the quiescence-promoting Forkhead Box O1 (*FOXO1*) gene, thus inducing its expression. The formation of this triplex involves an *Alu* element within the *MIRIAL* transcript, representing a previously undescribed mechanism of action for a lncRNA. Further, we generated a global *Mirial* knockout mouse line of. Angiogenic sprouting of aortic rings from *Mirial* knockout mice was reduced under basal conditions, but increased after VEGF administration, validating the *in vitro* angiogenic phenotype. Importantly, cardiac contractile function after acute myocardial infarction is severely reduced in *Mirial* knockout mice, as compared to wild-type littermates.

**Conclusions:** The lncRNA *MIRIAL* is an aging-induced regulator of endothelial quiescence and metabolism.

**Translational Perspective:** LncRNAs often exhibit cell-type or tissue-specific expression and regulation, rendering them potentially druggable targets requiring lower doses and having fewer side effects compared to protein targets. Our current research highlights, that loss of *Mirial* correlates with adverse outcomes post-acute myocardial infarction in a murine model. Dysregulation of *MIRIAL* in various human pathological conditions, such as ischemic heart disease, abdominal aortic aneurysm, cancer, and aging, indicates its potential as a diagnostic marker. Mechanistically, *MIRIAL* regulates endothelial quiescence by modulating *FOXO1* expression, suggesting it as a promising therapeutic target to counteract the age-related decline in endothelial cell function.

## 1. Introduction

LncRNAs are a heterogeneous class of RNA defined by a length exceeding 200 nucleotides and often exhibit a low abundance per cell. They can interact with DNA, other RNAs or proteins, thereby regulating gene expression, as well as RNA and protein function^1, 2^. Cardiovascular diseases (CVD) remain the leading cause of mortality worldwide^3^. The predominant risk factor for the development of CVD is aging^4^. In the vasculature, aging manifests as a gradual decline of endothelial function, stiffening of the vessel wall, compromised angiogenic capacity and chronic inflammation^5^. The transcription factor FOXO1 promotes endothelial quiescence and is indispensable for endothelial cell homeostasis, vasculogenesis, and development^6, 7^.

We identified an aging-induced lncRNA in murine cardiac endothelial cells that we termed *Mirial* (**Mi**cro**R**NA-cluster 23a∼27a∼24-2-associated and **I**nduced by **A**ging **L**ong non-coding RNA) (transcript ID: Gm26532, ENSMUSG00000097296), which is previously uncharacterized in endothelial cells. Prior publications have linked lower *Mirial* expression to reduced FoxO signaling in mice^8^. In humans, upregulation of the orthologous lncRNA *MIRIAL* (transcript IDs: LOC284454, AL832183, MIR23AHG, ENSG00000267519 or AC020916.1) was reported in nasopharyngeal carcinoma^9^ and hepatocellular carcinoma^10^, and was associated with elevated migration and invasion and poorer clinical prognosis. Furthermore, copy number variations of *MIRIAL* in colorectal carcinoma patient samples were described^11^. A study by Das *et al.* revealed that *MIRIAL* is a polyadenylated, single-exonic lncRNA with a half-life of 3 hours that is associated with chromatin. In contrast to the aforementioned studies of *MIRIAL* in cancer, they proposed its expression to be reduced in various types of cancers and cancer cell lines^12^. Lastly, *MIRIAL* was demonstrated to be upregulated and may serve as diagnostic biomarker in head and neck cancers^13^.

Here, we present a comprehensive characterization of the aging-induced lncRNA *MIRIAL* in endothelial cells, where it regulates metabolism and mitochondrial copy number by activating FOXO1 signaling. FOXO1 represses MYC signaling and promotes endothelial quiescence. Concurrent suppression of the p53 signaling pathway leads to enhanced proliferation and angiogenesis. Mechanistically, we discovered that an *Alu* element within the *MIRIAL* transcript forms an *RNA*·DNA:DNA triplex with a regulatory region of the *FOXO1* gene, resulting in its increased expression. Importantly, *MIRIAL* is the first lncRNA harboring a triplex-forming *Alu* element. Lastly, we generated a *Mirial* knockout mouse line, which presents with a sensitization of endothelial cells towards VEGF-A signaling in *ex vivo* experiments and a reduced myocardial contractile function after acute myocardial infarction (AMI) *in vivo*.

## 2. Materials and Methods

### 2.1. Animals

All mice experiments were carried out in accordance with the principles of laboratory animal care as well as according to the German national laws and the guidelines from Directive 2010/63/EU of the European Parliament on the protection of animals used for scientific purposes. The studies have been approved by the local ethics committee (Regierungspräsidium Darmstadt, Hesse, Germany). Mice were kept in individually ventilated cages (Tecniplast) at 12:12h-light/dark cycles at 21–24°C and 45-60% humidity. Water and ssniff R/M-H complete feed (ssniff Spezialdiäten) were fed *ad libitum*. Young (12 weeks) and aged (72 weeks) wild type C57Bl/6 mice were purchased from Janvier. The *MIRIAL* knockout mouse line was produced on a C57Bl/6N background by Cyagen/Taconic using CRISPR/Cas9. Both, male and female animals were used for the experiments. Animals were sacrificed by cervical dislocation under isoflurane (AbbVie) inhalation anesthesia. After perfusion with HBSS (Gibco), organs and tissues harvested for RNA or DNA isolation were snap-frozen in liquid nitrogen and homogenized in QIAzol lysis reagent (QIAGEN) or genomic lysis buffer (Invitrogen), respectively, using the Bead Mill MAX Homogeniser (VWR). The steps of RNA and genomic DNA isolation can be found elsewhere in this manuscript. AMI surgery was performed on 11–13-week-old animals by ligation of the left anterior descending (LAD) coronary artery as described before^14^. Inhalation anesthesia was induced using isoflurane (Abbvie). Analgesia was conducted using buprenorphine (0.1 mg/kg) injection and bupivacaine (0.25%, 1 mg/kg) infiltration. Post-operatively, buprenorphine (0.1 mg/kg) was injected twice a day for two days, and carprofen (5 mg/kg) once a day for six days. The animals’ health status was assessed daily. Four weeks after AMI surgery, mice were euthanized and perfused as described earlier and the hearts were harvested.

### 2.2. Cell culture

Pooled HUVECs were purchased from Lonza and cultured in Endothelial Cell Basal Medium (EBM) (Lonza) containing EGM SingleQuots supplements (Lonza) (without ascorbic acid) and 10% fetal bovine serum (FBS) (Gibco). HUVECs were used at passage 2 – 3 for experiments and cell number was determined using the Nucleocounter NC-2000 (Chemometec A/S). Human embryonic kidney (HEK) cell lines (Lenti-X 293T) (Takara) were cultured in high-glucose DMEM (Gibco) with 10 % FBS (Gibco) and 1x Pen/Strep (Roche). Lenti-X 293T cells were kindly provided by the lab of Ralf Brandes/Matthias Leisegang. All cells were regularly tested for mycoplasma contamination and kept at 37°C and 5% CO_2_ in a humidified atmosphere.

### 2.3. Human samples

Participants gave informed written consent prior to the inclusion in the study. The investigation conformed to the Declaration of Helsinki. Left ventricular tissue samples from ischemic heart disease (ISHD) patients and control samples were sourced from the University of Sydney (Sydney, NSW, Australia) with approval from the Human Research Ethics Committee (number 2012/2814). Control samples were obtained from explanted hearts of heathy donors who died from non-cardiac causes, typically motor vehicle accidents.

### 2.4. RNA-sequencing

HUVECs were transfected with either LNA Control or LNA MIRIAL. 48 h post-transfection, RNA was isolated as described earlier. RNA quality control, library preparation, RNA-sequencing and analysis was performed as described in^15^. The related data are available on Gene Expression Omnibus (accession number GSE248795; reviewer token: cxutusselbenlkp; https://www.ncbi.nlm.nih.gov/geo/query/acc.cgi?acc=GSE248795). Single-cell RNA-sequencing data of human aneurysmal abdominal aortas was obtained from Gene Expression Omnibus (accession number GSE237230)^16^. RNA-sequencing data of cardiac endothelial cells derived from young and aged mice was published previously and is available online^17^.

### 2.5. Statistics

Statistical analysis was performed using GraphPad Prism 7.02, 8.0.1 or 9.0.2 for Windows (GraphPad Software, www.graphpad.com). Results are presented as mean ± standard deviation (SD) or standard error of the mean (SEM). Gaussian distribution was assessed using the Shapiro-Wilk normality test. To compare two conditions, paired or unpaired student’s *t* test, Mann-Whitney test or Kolmogorov-Smirnov tests were used. To compare multiple conditions, we used (Brown-Forsythe and Welch) analysis of variance (ANOVA) or Kruskal-Wallis test with Tukey *post-hoc* test.

## 3. Results

### 3.1. *MIRIAL* is an aging-induced long non-coding RNA that regulates endothelial cell quiescence

*Mirial* (transcript ID: Gm26532) was first identified in an RNA-sequencing data set generated from cardiac endothelial cells of 12 weeks old and 20 months old mice **(Fig. 1A)** ^17^. These data showed that *Mirial* was upregulated during aging, which was further confirmed in aortic intima samples of young (12 weeks) and aged (20 months) mice analyzed by qRT-PCR **(Fig. 1B)** . The *Mirial* gene locus also encompasses the miRNA cluster 23a∼27a∼24-2 and is highly conserved. By analyzing the corresponding locus in humans, the human orthologue of *Mirial* could be identified **(Supp. Fig. 1A)** . Interestingly, *MIRIAL* was also found to be upregulated in an RNA-sequencing data set of human native endothelial cells from young and aged individuals^18^. Previously, it had been shown that *MIRIAL* is highly sequence conserved solely in primates^12^. To assess the degree of sequence conservation between human and mouse we employed the LALIGN DNA:DNA tool^19^, which revealed that around 50% of the mouse sequence is also present in the human orthologue **(Supp. Fig. 1B)** . Comparing the coding probability of *MIRIAL* with known transcripts using the Coding-Potential Assessment Tool (CPAT)^20^ we found that *MIRIAL* exhibited a coding probability similar to well-known lncRNAs such as *XIST* and *MALAT1*. Conversely, protein-coding transcripts like *GAPDH*, but also *ZSWIM4*, the protein-coding gene upstream in the locus of *MIRIAL*, were found on the opposite side of the spectrum **(Supp. Fig. 1C)** . By performing RNA-sequencing analyses, we assessed the transcription levels of *MIRIAL* in various human cell types of cardiovascular origin. The results demonstrated that *MIRIAL* is expressed in endothelial cells of different vascular beds, as well as in other non-endothelial cell types including cardiomyocytes **(Supp. Fig. 1D)** . In the context of CVDs, *MIRIAL* expression was induced in left ventricular tissue samples from patients with ischemic heart disease (ISHD) **(Fig. 1C)** and in endothelial cells from human abdominal aortic aneurysm (AAA) samples using single-cell RNA-sequencing **(Supp. Fig. 1E)**^16^. To investigate the function of *MIRIAL*, we chose a loss-of-function approach using LNA gapmers in human umbilical vein endothelial cells (HUVECs), achieving a knockdown efficiency of approximately 80-88% **(Supp. Fig. 2A)** . *MIRIAL* silencing resulted in a decrease in proliferation **(Fig. 1D)** with a block of the cell cycle at the G0/G1 phase **(Fig. 1E)**. A scratch-wound assay combined with electric cell-substrate impedance sensing (ECIS) showed reduced barrier function **(Supp. Fig. 2B)** and impaired migration **(Fig. 1F)** after *MIRIAL* knockdown. Next, we assessed whether these effects were due to quiescence (a reversible reduction in cellular activity), senescence or apoptosis (both of which are irreversible). Apoptosis was unchanged in *MIRIAL* knockdown cells, as assessed by Caspase-3/-7 activity measurement **(Fig. 1G)** . In addition, there was no change in number of senescent cells after *MIRIAL* knockdown using a senescence-associated β-galactosidase staining **(Supp. Fig. 2C)**, DNA-damage-associated y-H2A.X staining **(Supp. Fig. 2D)** or telomere length measured by qPCR **(Supp. Fig. 2E)** . These results suggested that loss of *MIRIAL* induces quiescence rather than senescence or apoptosis. Given the observed decrease in proliferation and migration, we hypothesized that angiogenic sprouting would also be reduced. This was confirmed using a spheroid sprouting assay under unstimulated conditions. Importantly, stimulation with VEGF-A resulted in an increase in angiogenic sprouting when *MIRIAL* was silenced, as compared to VEGF-A stimulated control cells **(Fig. 1H)**. Moreover, VEGF-A stimulation rescued the effects of *MIRIAL* knockdown on proliferation **(Supp. Fig. 2F)** . Taken together, these results suggest that loss of *MIRIAL* induces a reversible state of quiescence of endothelial cells.

**Fig. 1:**
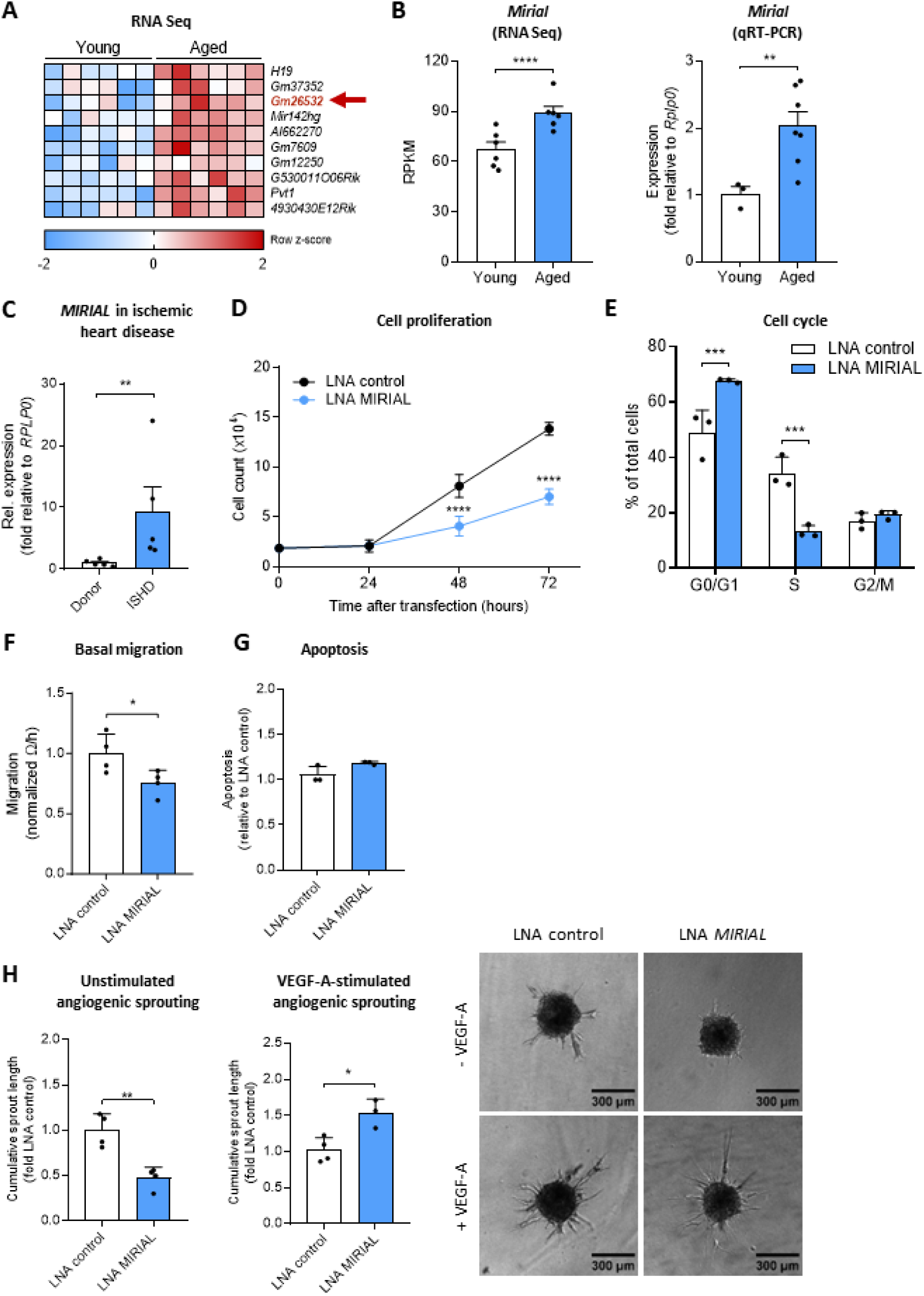
*MIRIAL* is an aging-induced long non-coding RNA that regulates endothelial cell quiescence. **A** Heatmap representing the most significant differentially regulated lncRNAs from RNA-sequencing data of cardiac ECs isolated from young (12 weeks) and aged (20 months) mice (*n*=6). **B** *Mirial* expression from RNA seq data (*n*=6; Mann-Whitney test with FDR correction) quantified from fig. 1 A and *Mirial* expression from aortic intima of young (12 weeks) and aged mice (20 months) measured by qRT-PCR (n=3 and 7; Welch’s *t* test). Results are expressed as mean ± s.e.m. **C** *MIRIAL* expression in ischemic heart disease cardiac biopsies compared to healthy controls measured by qRT-PCR (*n*=5; Mann-Whitney test). Results are expressed as mean ± s.e.m. **D** Growth curve depicting the cell number of HUVECs after the indicated hours (0-72h) of gapmeR-mediated *MIRIAL* knockdown (20 nM) compared to control-transfected HUVECs (representative graph of *n*=3; ANOVA). **E** EdU combined with FxCycle flow cytometry-based cell cycle analysis of HUVECs 48h after gapmeR-mediated *MIRIAL* knockdown (20 nM) compared to control-transfected HUVECs (*n*=3; ANOVA). **F** Electric cell-substrate impedance sensing (ECIS)-based migration assay of HUVECs 48h after gapmeR-mediated *MIRIAL* knockdown (20 nM) compared to control-transfected HUVECs (*n*=4; unpaired *t* test). **G** Caspase 3/7 activity-based apoptosis assay of HUVECs 48h after gapmeR-mediated *MIRIAL* knockdown (20 nM) compared to control-transfected HUVECs (*n*=3; unpaired *t* test). **H** Cumulative sprout length in an angiogenic spheroid sprouting assay under basal and VEGF-stimulated (50 ng/mL) conditions using HUVECs 72h after gapmeR-mediated *MIRIAL* knockdown (20 nM) compared to control-transfected HUVECs (*n*=4; unpaired *t* test). 8-15 spheroids per condition were analyzed. Results are expressed as mean ± s.d. unless otherwise stated; \**p*<0.05, \*\**p*<0.01, \*\*\**p*<0.001, **** *p*<0.0001

### 3.2. *MIRIAL* regulates endothelial mitochondrial metabolism

Glycolysis serves as the predominant pathway for energy production in endothelial cells and plays a central role in endothelial quiescence^21^. To investigate the potential impact of *MIRIAL* on energy homeostasis in endothelial cells, we used the Seahorse glycolysis stress test. Surprisingly, *MIRIAL* knockdown did not affect glycolysis (**Supp. Fig. 3A** and **3B)** . However, using the mitochondrial stress test, we found that *MIRIAL* knockdown led to an increase in maximal respiration **(Fig. 2A** and **2B)** and spare respiratory capacity (SRC) **(Fig. 2C)**, while basal respiration remained unchanged **(Fig. 2A** and **2B)** . Furthermore, ATP production **(Fig. 2D)** and cellular ATP levels were increased **(Fig. 2E)** after *MIRIAL* knockdown. To determine the underlying cause of this apparent increase in mitochondrial activity, we analyzed mitochondrial content after knockdown of *MIRIAL*. We found an increase in mitochondrial mass **(Fig. 2F)** and mitochondrial DNA content **(Fig. 2G)** relative to nuclear DNA following *MIRIAL* knockdown. Nevertheless, mitochondrial membrane potential **(Supp. Fig. 3C)** and production of reactive oxygen species (ROS) **(Supp. Fig. 3D)** remained unchanged, suggesting that *MIRIAL* silencing induces the number of functional mitochondria.

**Fig. 2:**
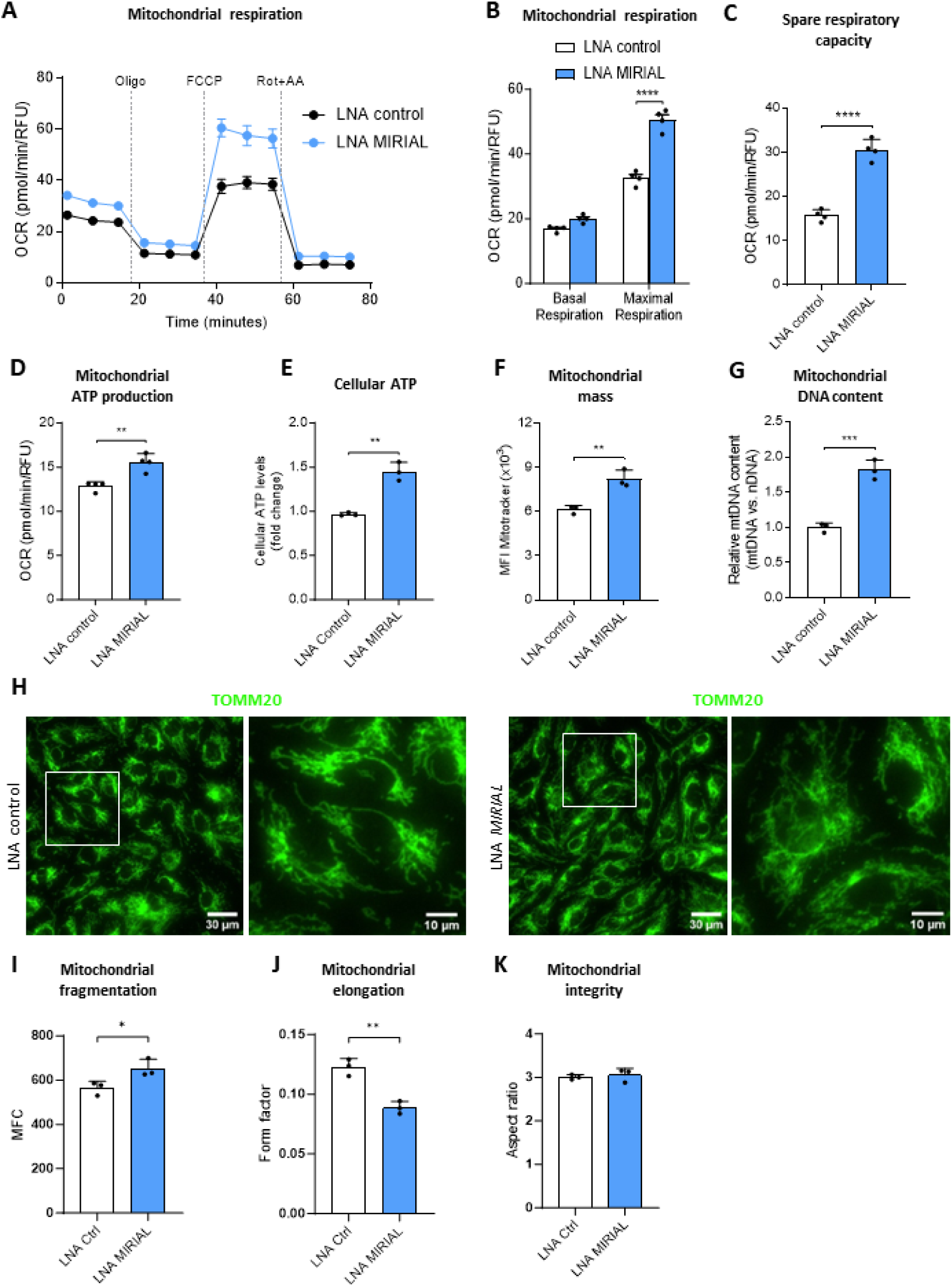
*MIRIAL* regulates endothelial mitochondrial metabolism. **A** Oxygen consumption rate (OCR) as a measure of mitochondrial respiration in HUVECs 48h after gapmeR-mediated *MIRIAL* knockdown (20 nM) compared to control-transfected HUVECs using Seahorse mitochondrial stress test (representative graph of *n*=4). **B** Oxygen consumption rate (OCR) as a measure of mitochondrial basal and maximal respiration in HUVECs 48h after gapmeR-mediated *MIRIAL* knockdown (20 nM) compared to control-transfected HUVECs, quantified from fig. 2 A using Seahorse mitochondrial stress test (*n*=4; ANOVA). **C** Spare respiratory capacity (SRC) of HUVECs 48h after gapmeR-mediated *MIRIAL* knockdown (20 nM) compared to control-transfected HUVECs, quantification from fig. 4A (*n*=4; unpaired *t* test). **D** Mitochondrial ATP production of HUVECs 48h after gapmeR-mediated *MIRIAL* knockdown (20 nM) compared to control-transfected HUVECs using Seahorse mitochondrial stress test (*n*=4; unpaired *t* test). **E** Cellular ATP levels of HUVECs 48h after gapmeR-mediated *MIRIAL* knockdown (20 nM) compared to control-transfected HUVECs using bioluminescence-based assay (*n*=3). **F** Mitochondrial mass of HUVECs 48h after gapmeR-mediated *MIRIAL* knockdown (20 nM) compared to control-transfected HUVECs measured by Mitotracker flow cytometry (*n*=3; unpaired *t* test). **G** Mitochondrial DNA (mtDNA) content relative to nuclear DNA (nDNA) content of HUVECs 48h after gapmeR-mediated *MIRIAL* knockdown (20 nM) compared to control-transfected HUVECs using qPCR (*n*=3; unpaired *t* test). **H** Representative images of HUVECs 48h after gapmeR-mediated *MIRIAL* knockdown (20 nM) compared to control-transfected HUVECs stained with an TOMM20 antibody and analyzed by immunofluorescence microscopy. **I** Mitochondrial fragmentation count (MFC) in HUVECs 48h after gapmeR-mediated *MIRIAL* knockdown (20 nM) compared to control-transfected HUVECs stained with an TOMM20 antibody and analyzed by immunofluorescence microscopy (*n*=3; unpaired *t* test). **J** Mitochondrial elongation measured as mitochondrial form factor in HUVECs 48h after gapmeR-mediated *MIRIAL* knockdown (20 nM) compared to control-transfected HUVECs stained with an TOMM20 antibody and analyzed by immunofluorescence microscopy (*n*=3; unpaired *t* test). **K** Mitochondrial integrity in HUVECs 48h after gapmeR-mediated *MIRIAL* knockdown (20 nM) compared to control-transfected HUVECs stained with an TOMM20 antibody and analyzed by immunofluorescence microscopy (*n*=3; unpaired *t* test). Results are expressed as mean ± s.d.; \**p*<0.05, \*\**p*<0.01, \*\*\**p*<0.001, **** *p*<0.0001

The tightly regulated processes of fission and fusion play pivotal roles in maintaining the cellular oxidative capacity, energy supply and biomass supply through coordinating mitochondrial copy number^22^. Accordingly, we wanted to explore whether these processes are involved in the mechanism by which *MIRIAL* knockdown augments mitochondrial copy number. Consistent with the increase of total mitochondrial mass and mitochondrial DNA content after knockdown of *MIRIAL*, we also found an elevation in the expression levels of almost all key players relevant for fission and fusion, including mitochondrial fission factor (MFF), dynamin 1-like protein (DRP1), Mitofusin-1, Mitofusin-2, mitochondrial dynamin like GTPase Optic Atrophy 1 (OPA1) **(Supp. Fig. 4 A-G)** and the transporter of the outer mitochondrial membrane (TOMM20) **(Supp. Fig. 4H)**, responsible for mitochondrial membrane transport. To investigate the impact of increased levels of fission and fusion proteins on mitochondrial morphology following *MIRIAL* knockdown, we utilized TOMM20 staining followed by immunofluorescence microscopy **(Fig. 2H)** . We quantified the mitochondrial fragmentation count (MFC), aspect ratio (a measure of mitochondrial length and structural integrity), and form factor (a combined measure of mitochondrial elongation and branching). Our results showed a significant increase in the fragmentation count **(Fig. 2I)** and a reduction in the form factor **(Fig. 2J)** upon *MIRIAL* knockdown. However, the aspect ratio remained unchanged **(Fig. 2K)**. These findings suggest that the loss of *MIRIAL* increases mitochondrial content and function, but this effect does not seem to be mediated through changes in fission or fusion. The increased fragmentation count observed after *MIRIAL* silencing indicates a higher number of mitochondria. These mitochondria exhibit less branching, yet their individual structural integrity and length remain unaltered. In summary, these results show that knockdown of *MIRIAL* heightens functional mitochondrial content in endothelial cells.

### 3.3. *MIRIAL* knockdown in HUVECs activates the p53 signaling pathway

To gain insight into the role of *MIRIAL* in endothelial cell quiescence, we first focused on its subcellular localization. Subcellular fractionation of HUVECs into cytoplasmic, nucleoplasmic and chromatin fractions revealed that *MIRIAL* was predominantly present in the nucleus (∼80%), with a significant proportion present in the chromatin fraction **(Supp. Fig. 4I)**. Given this localization pattern, we hypothesized that *MIRIAL* might directly regulate gene expression. To assess its impact on gene expression we performed RNA-sequencing of HUVECs after *MIRIAL* knockdown, comparing them to control-transfected cells. Following *MIRIAL* knockdown, 974 genes were differentially expressed (Adjusted *p*-value < 0.01, absolute log_2_(fold change) > 0.585) **(Fig. 3A)** . Kyoto Encyclopedia of Genes and Genomes (KEGG) pathway enrichment analysis demonstrated that genes associated with the cell cycle were downregulated significantly **(Fig. 3B)**, aligning with our previous findings on cell proliferation **(Fig. 1C)** . In addition, we observed a significant upregulation in genes annotated as members of the p53 signaling pathway **(Fig. 3B)** .

**Fig. 3:**
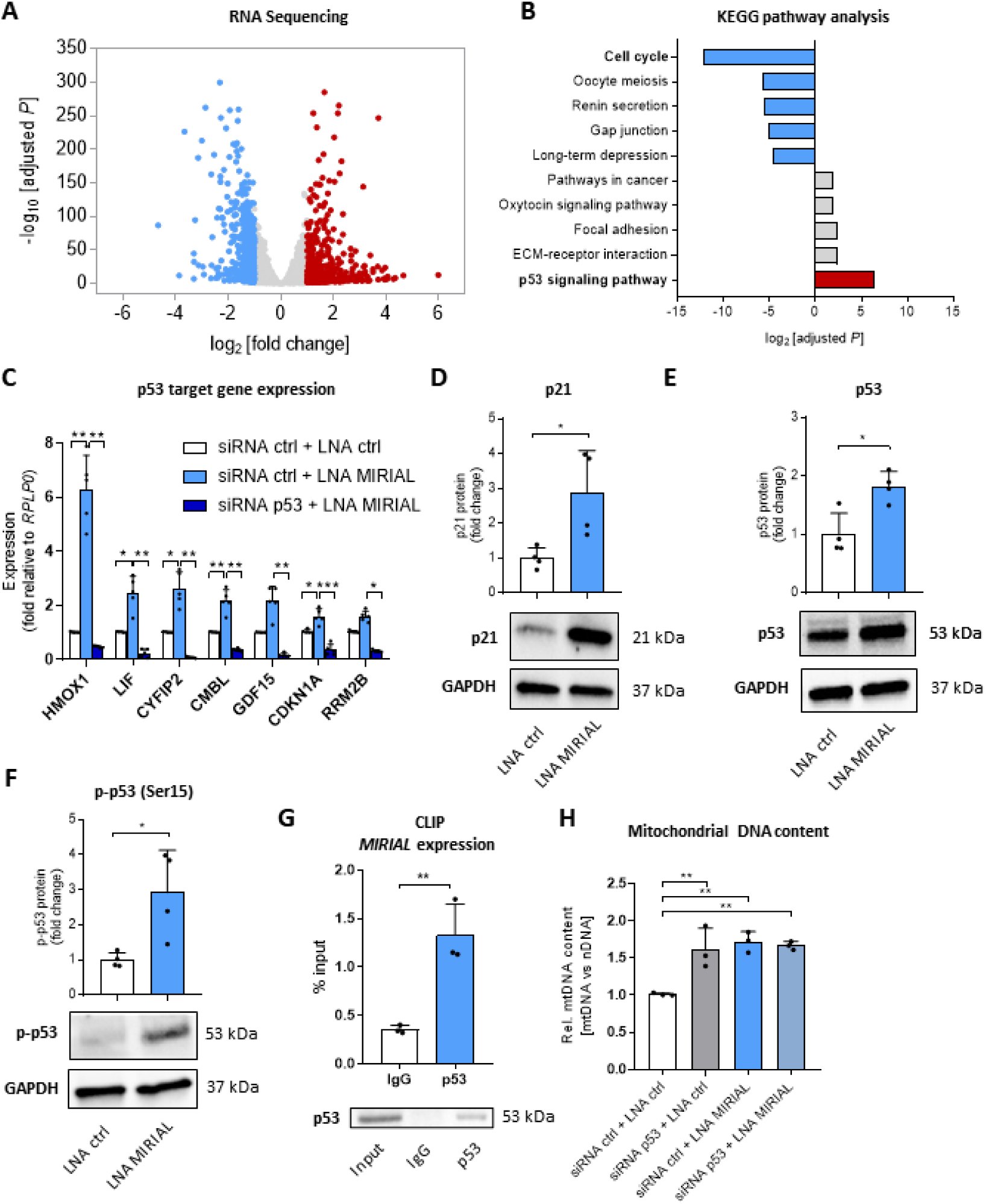
*MIRIAL* knockdown in HUVECs activates the p53 signaling pathway. **A** Volcano plot representing differentially expressed genes from RNA-sequencing of HUVECs 48h after transfection with either gapmeRs targeting *MIRIAL* (20 nM) or control gapmeRs (control vs *MIRIAL* knockdown; colored dots fold change≥2; adj. *p*≤0.01.; *n*=4). **B** KEGG pathway analysis of RNA-sequencing data in fig. 3A. **C** p53 target gene expression in HUVECs 48h after gapmeR-mediated *MIRIAL* knockdown (20 nM) compared to control-transfected HUVECs and siRNA-mediated knockdown of p53 (10 nM) or control transfected HUVECs, using qRT-PCR (*n*=5; Brown-Forsythe and Welch ANOVA). **D** Protein levels of p21 in HUVECs 48h after gapmeR-mediated *MIRIAL* knockdown (20 nM) compared to control-transfected HUVECs using Western Blot (*n*=4; unpaired *t* test). **E** Protein levels of p53 in HUVECs 48h after gapmeR-mediated *MIRIAL* knockdown (20 nM) compared to control-transfected HUVECs using Western Blot (*n*=4; unpaired *t* test). **F** Protein levels of phosphorylated p53 (Ser15) in HUVECs 48h after gapmeR-mediated *MIRIAL* knockdown (20 nM) compared to control-transfected HUVECs using Western Blot (*n*=4; unpaired *t* test). **G** *MIRIAL* expression measured by qRT-PCR after crosslinking and RNA immunoprecipitation (CLIP) of HUVEC lysate with a p53 antibody or IgG isotype control antibody (5 µg each) and Western blot analysis of p53 in the input and immunoprecipitates (*n*=3; unpaired *t* test). **H** Mitochondrial DNA (mtDNA) content relative to nuclear DNA (nDNA) content of HUVECs 48h after gapmeR-mediated *MIRIAL* knockdown (20 nM) compared to control-transfected HUVECs and siRNA-mediated knockdown of p53 (10 nM) or control transfected HUVECs using qPCR (*n*=3; ANOVA). Results are expressed as mean ± s.d.; \**p*<0.05, \*\**p*<0.01, \*\*\**p*<0.001

Under basal conditions, p53 is constantly expressed and subsequently poly-ubiquitinylated by the E3 ubiquitin-protein ligase MDM2, leading to its proteasomal degradation. However, in response to various stressors, such as metabolic or oxidative stress, p53 undergoes phosphorylation at multiple sites, resulting in its stabilization and nuclear translocation. In the nucleus, p53 acts as a transcription factor regulating gene expression^23^. We validated the activation of p53 target genes using qRT-PCR, showing that *MIRIAL* knockdown indeed induces expression of typical p53 target genes (*HMOX1*, *LIF*, *CYFIP2*, *CMBL*, *GDF15*, *CDKN1A*, and *RRM2B*). Importantly, the upregulation of p53 target genes was fully dependent on the presence of p53, since knockdown of p53 using siRNA, rescued the loss-of-*MIRIAL*-mediated induction of these target genes **(Fig. 3C)**. In addition, we confirmed the upregulation of p21 (*CDKN1A*) **(Fig. 3D)**, p53 **(Fig. 3E)** and phosphorylation of p53 at Ser15 **(Fig. 3F)** on the protein level and discovered a physical interaction between p53 and *MIRIAL* using crosslinking and RNA immunoprecipitation (CLIP) **(Fig. 3G)** . Interestingly, the increase in mitochondrial DNA in *MIRIAL* knockdown cells could not be rescued by a simultaneous knockdown of p53 **(Fig. 3H)**, indicating that p53 induction is not responsible for the observed increase in mitochondrial activity following *MIRIAL* silencing.

The primary transcript of the *MIRIAL* locus which contains both the miRNAs and *MIRIAL* is immediately cleaved by Drosha after transcription^12^. Given the mutual transcription of the miRNAs and *MIRIAL*, it was essential to investigate whether a potential decrease in miRNA abundance contributed to the phenotype. Using TaqMan miRNA assay probes, we assessed the expression levels of all three members of the miRNA cluster (miR-23a, miR-27a and miR-24-2) after *MIRIAL* knockdown. All three miRNAs were downregulated following *MIRIAL* knockdown **(Supp. Fig. 5A)**. If this reduction were functionally relevant, we would expect an upregulation of the target genes regulated by these miRNAs. However, expression levels of two selected target genes of each miRNA did not increase after *MIRIAL* knockdown **(Supp. Fig. 5B)**, suggesting that the moderate reduction in miRNA expression did not result in target de-repression. Nonetheless, to exclude the possibility that the observed phenotype resulting from silencing of *MIRIAL* is due to reduction of these miRNAs, we performed miRNA overexpression experiments by transfecting HUVECs with combined pre-miRNAs achieving expression levels 40-150-fold higher than control **(Supp. Fig. 5C)**. Notably, *MIRIAL* knockdown efficiency was not affected by miRNA overexpression **(Supp. Fig. 5D)** . We further assessed whether miRNA overexpression could rescue the functional effects of *MIRIAL* knockdown on cell proliferation, mitochondrial DNA content and p53 target gene expression. While miRNA overexpression increased proliferation, this effect was independent of *MIRIAL*, as knockdown of *MIRIAL* still significantly decreased cell numbers **(Supp. Fig. 5E)** . Additionally, mitochondrial DNA content remained elevated after *MIRIAL* knockdown, despite overexpression of the miRNAs **(Supp. Fig. 5F)** . Regarding p53 target gene expression, we observed that *HMOX1*, *RRM2B* and *MDM2* were the only genes influenced by miRNA overexpression, with *HMOX1* being a known target of miR-24^24^ **(Supp. Fig. 5G)**. These findings strongly suggest that the effects of *MIRIAL* silencing on endothelial cells are unlikely to be solely attributed to a reduction in miR-23a, miR-27a and miR-24-2 expression.

In conclusion, the results from these loss-of-function experiments provide evidence that *MIRIAL* acts as a repressor of p53 signaling and interacts with p53. However, this pathway does not contribute to the *MIRIAL*-mediated reduction of mitochondrial content. furthermore, the effects of *MIRIAL* silencing on endothelial quiescence, mitochondrial activity and p53 signaling are independent of the co-expressed miR-23a, miR-27a and miR-24-2.

### 3.4. *MIRIAL* regulates endothelial quiescence through FOXO1 and MYC signaling

To uncover the underlying mechanism by which *MIRIAL* regulates mitochondrial biogenesis, we investigated the involvement of key transcription factors that govern endothelial metabolism and quiescence, namely FOXO1 and MYC (also known as c-myc). FOXO1 and MYC exhibit antagonistic roles in the endothelium. Under conditions lacking growth factor stimulation, FOXO1 inhibits MYC signaling and promotes endothelial quiescence by decreasing proliferation and metabolism^6^. However, upon stimulation with growth factors, e.g. VEGF-A, this balance shifts towards enhanced MYC signaling. VEGF-A leads to an increase of Akt kinase activity, resulting in phosphorylation of FOXO1, subsequent nuclear export and proteasomal degradation^25^.

By comparing gene sets of FOXO1 target genes (obtained from GSEA Molecular Signatures Database C3 https://www.gsea-msigdb.org/gsea/msigdb/human/geneset/FOXO1_02.html) and MYC mitochondrial target genes (obtained from^26^) to expression of all genes after *MIRIAL* knockdown, we observed a robust downregulation (left-shift) of the FOXO1 target genes, and an overall upregulation (right-shift) of mitochondrial target genes encoded in the nucleus and controlled by MYC **(Fig. 4A)**.

**Fig. 4:**
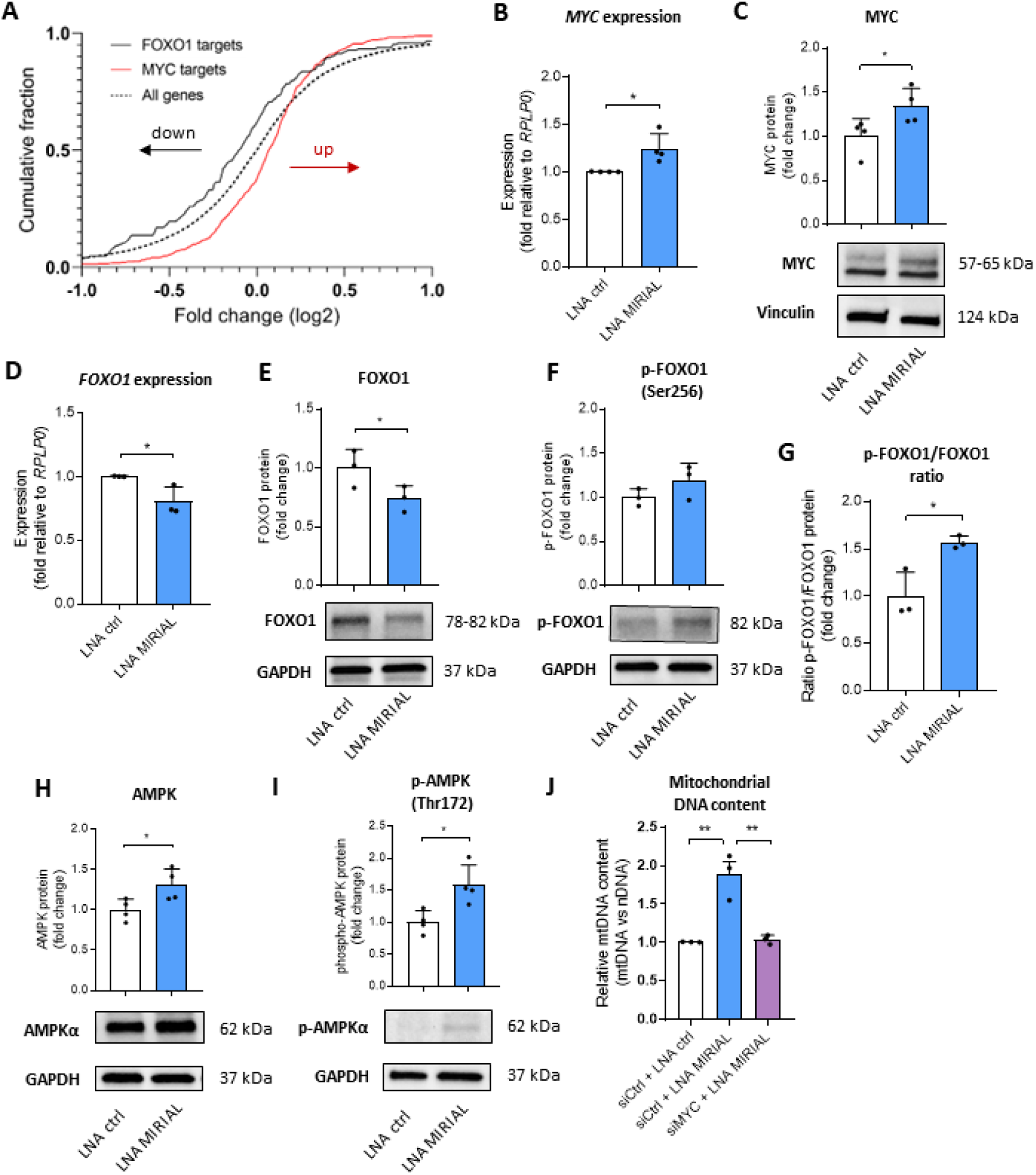
*MIRIAL* regulates endothelial quiescence through FOXO1 and MYC. **A** Expression changes of MYC nuclear target genes encoding mitochondrial proteins and FOXO1 target genes compared to all genes 48h after gapmeR-mediated *MIRIAL* knockdown (20 nM) in HUVECs compared to control-transfected HUVECs using RNAseq (*n*=4). **B** *MYC* expression in HUVECs 48h after gapmeR-mediated *MIRIAL* knockdown (20 nM) compared to control-transfected HUVECs using qPCR (*n*=4; Mann-Whitney test). **C** Protein levels of MYC in HUVECs 48h after gapmeR-mediated *MIRIAL* knockdown (20 nM) compared to control-transfected HUVECs using Western blot (*n*=4; Mann-Whitney test). **D** *FOXO1* expression in HUVECs 48h after gapmeR-mediated *MIRIAL* knockdown (20 nM) compared to control-transfected HUVECs using qRT-PCR (*n*=3; unpaired *t* test). **E** Protein levels of FOXO1 in HUVECs 48h after gapmeR-mediated *MIRIAL* knockdown (20 nM) compared to control-transfected HUVECs using Western blot (*n*=3; paired *t* test). **F** Protein levels of phosphorylated FOXO1 (Ser256) in HUVECs 48h after gapmeR-mediated *MIRIAL* knockdown (20 nM) compared to control-transfected HUVECs using Western blot (*n*=3; paired *t* test). **G** Ratio of phosphorylated to total protein levels of FOXO1 in HUVECs 48h after gapmeR-mediated *MIRIAL* knockdown (20 nM) compared to control-transfected HUVECs quantified from fig. 4 E and F (*n*=3; unpaired *t* test). **H** Protein levels of AMPK in HUVECs 48h after gapmeR-mediated *MIRIAL* knockdown (20 nM) compared to control-transfected HUVECs using Western blot (*n*=4; unpaired *t* test). **I** Protein levels of phosphorylated AMPK (Thr172) in HUVECs 48h after gapmeR-mediated *MIRIAL* knockdown (20 nM) compared to control-transfected HUVECs using Western blot (*n*=4; unpaired *t* test). **J** Mitochondrial DNA (mtDNA) content relative to nuclear DNA (nDNA) content of HUVECs 48h after gapmeR-mediated *MIRIAL* knockdown (20 nM) compared to control-transfected HUVECs and siRNA-mediated knockdown of MYC (60 nM) or control transfected HUVECs using qPCR (*n*=3; ANOVA). Results are expressed as mean ± s.d.; \**p*<0.05, \*\**p*<0.01

After *MIRIAL* knockdown, we observed an increase in *MYC* expression in RNA-sequencing **(Supp. Fig. 6A)**, qRT-PCR **(Fig. 4B)** and at protein level in Western blot **(Fig. 4C)** . Simultaneously, *FOXO1* expression was decreased in RNA-sequencing **(Supp. Fig. 6B)**, qRT-PCR **(Fig. 4D)** and at protein level **(Fig. 4E)**. Additionally, we found elevated levels of FOXO1 phosphorylation at Ser256 **(Fig. 4F)** and an increased ratio of phospho-FOXO1/total FOXO1 **(Fig. 4G)**. Both are indicative of inactivation, nuclear export, and subsequent degradation of FOXO1. Activation of the anabolic transcription factor MYC leads to the exhaustion of energy reserves, which in turn triggers the activation of AMP-activated kinase (AMPK), a key sensor of cellular energy status^27^. Under conditions of a high AMP:ATP ratio, activated AMPK can attenuate the cell cycle through p53 phosphorylation at Ser15 until energy stocks are replenished^28, 29^. We assessed whether AMPK activation links MYC to the induction of the p53 signaling pathway after *MIRIAL* knockdown using Western blot analysis. Protein levels of AMPK **(Fig. 4H)** and its activated, phosphorylated (Thr172) form **(Fig. 4I)** were both increased. MYC is a potent regulator of mitochondrial biogenesis and respiration^30, 31^. So, we hypothesized that the upregulation of MYC after *MIRIAL* silencing is responsible for the induction of mitochondrial function. To test this hypothesis, we performed siRNA-mediated knockdown of MYC in conjunction with *MIRIAL* knockdown. The resulting abrogation of *MIRIAL* depletion-mediated increase in mtDNA content demonstrated that MYC is indeed essential for the regulation of mitochondrial biogenesis by *MIRIAL* **(Fig. 4J)**.

### 3.5. *MIRIAL* forms *RNA*·DNA:DNA triplexes in the *FOXO1* promoter region

Certain lncRNAs can act as epigenetic modifiers in a sequence-specific manner by forming an *RNA*·DNA:DNA triplex with DNA. In this mechanism, a single-stranded lncRNA molecule inserts into the major groove of the DNA double helix *via* Hoogsteen base pairing. This structural arrangement enables the lncRNA to influence gene expression through chromatin remodeling and recruitment of epigenetic modifiers^32, 33^. Examples include the lncRNAs *Sarrah*^34^ and *HIF1α-AS1*^35^. In our study, we observed that *MIRIAL* is primarily associated with the chromatin in subcellular fractionation experiments **(Fig. 3A)** . Given its localization, we investigated whether *MIRIAL* can form triplexes with double-stranded DNA. We analyzed potential triplex-forming regions (TFRs) in *MIRIAL* using *TriplexAligner*^36^ and found one region within *MIRIAL* that has a high propensity for predicted triplex formation with double-stranded DNA **(Fig. 5A)** . *MIRIAL* has previously been described as containing four transposable elements that belong to the class of short, interspersed elements (SINEs), more precisely, *Alu* elements^12^. The TFR we identified corresponds to the localization of the first *Alu* element. Transposable elements, in particular *Alu* elements, have been proposed as functional regions of lncRNAs before, e.g. for the lncRNAs S*MANTIS (* formerly known as *MANTIS)*^37, 38^ and *ANRIL*^39^.

**Fig. 5:**
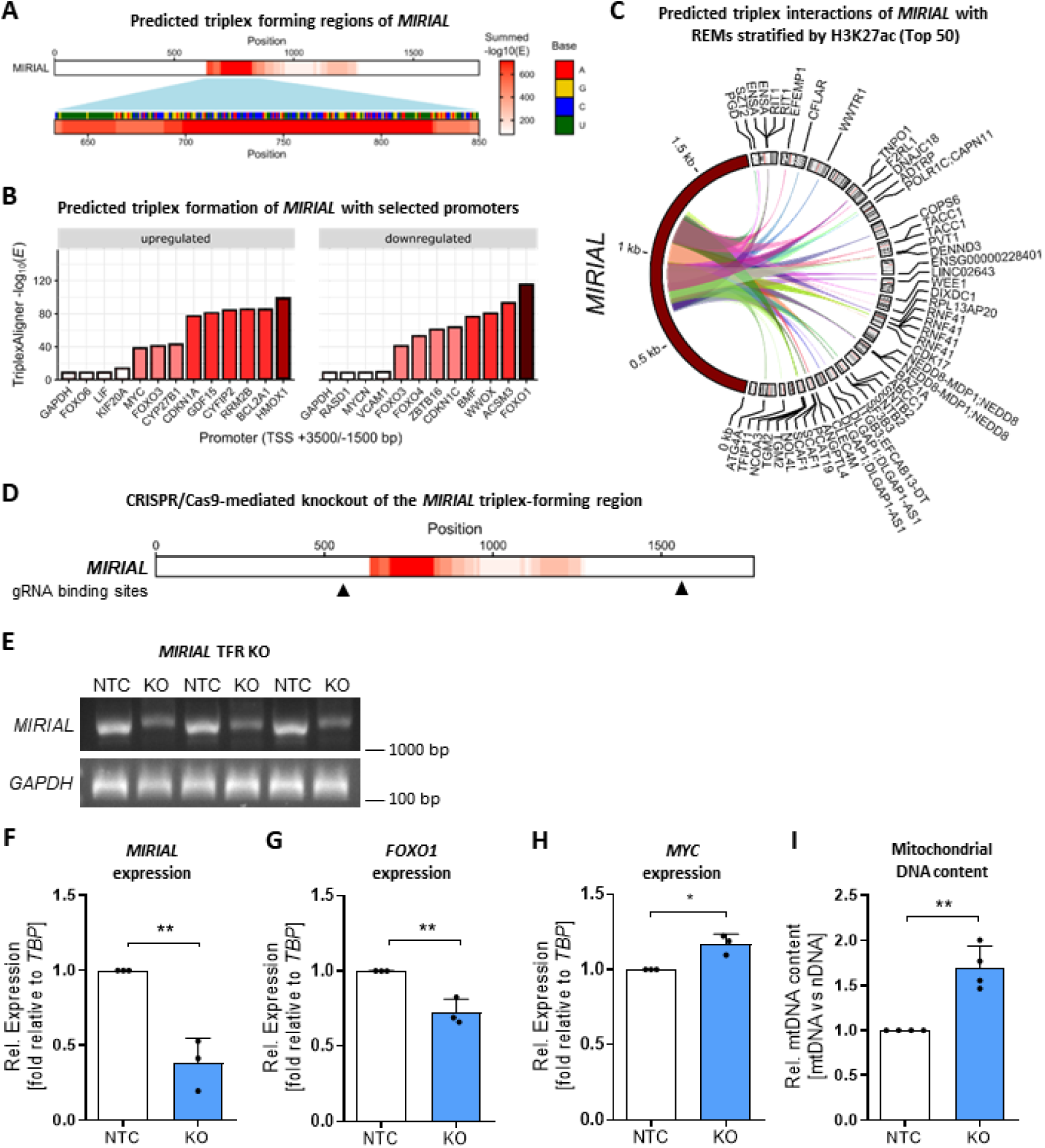
*MIRIAL* forms *RNA*·DNA:DNA triplexes in the *FOXO1* promoter region. **A** Schematic representation of the lncRNA *MIRIAL* showing predicted probability (-log10(E) values) of *RNA***·**DNA:DNA triplex forming regions (TFRs) in the lncRNA using *TriplexAligner***. B** *TriplexAligner* scores (-log10(E) values) for predicted interactions between the lncRNA *MIRIAL* and selected promoters of genes upregulated after *MIRIAL* knockdown (*FOXO6, LIF, KIF20A, MYC, CYP27B1, CDKN1A, GDF15, CYFIP2, RRM2B, BCL2A1, HMOX1*) or downregulated after *MIRIAL* knockdown (*RASD1, MYCN, VCAM1, FOXO4, ZBTB16, CDKN1C, BMF, WWOX, ACSM3, FOXO1*) in RNAseq (fig. 3A), in comparison to negative control promoters of genes that are not regulated after *MIRIAL* knockdown (*GAPDH* and *FOX O*)*3* using *TriplexAligner*. **C** Top 50 predicted triplex interactions of *MIRIAL* with regulatory elements (REMs) using *TriplexAligner* ranked by *TriplexAligner* score and stratified by genome accessibility (H3K27ac) data obtained from ENCODE. **D** Schematic representation of the *MIRIAL* transcript and the binding sites of guide RNAs (gRNAs) used for CRISPR/Cas9-mediated knockout (KO) of the triplex-forming region (TFR) of *MIRIAL*. **E** CRISPR/Cas9-mediated knockout (KO) of the *MIRIAL* TFR in HUVECs using lentiviral transduction of *MIRIAL*-targeting guide-RNAs or non-targeting control (NTC) gRNAs and measured my genomic DNA PCR 10 days post-transduction. *GAPDH* serves as loading control. Three independent experiments are shown. **F** *MIRIAL* expression after CRISPR/Cas9-mediated knockout (KO) of the *MIRIAL* TFR or non-targeting control (NTC) using qRT-PCR (*n*=3; unpaired *t* test). **G** *FOXO1* expression in HUVECs 10 days after CRISPR/Cas9-mediated *MIRIAL* TFR knockout compared to control-transduced HUVECs using qRT-PCR (*n*=3; unpaired *t* test). **H** *MYC* expression in HUVECs 10 days after CRISPR/Cas9-mediated *MIRIAL* TFR knockout compared to control-transduced HUVECs using qRT-PCR (*n*=3; unpaired *t* test). **I** Mitochondrial DNA (mtDNA) content relative to nuclear DNA (nDNA) content of HUVECs 10 days after CRISPR/Cas9-mediated *MIRIAL* TFR knockout compared to control-transduced HUVECs using qPCR (*n*=4; unpaired *t* test). Results are expressed as mean ± s.d.; \**p*<0.05, \*\**p*<0.01

Further, we were interested to identify potential triplex target sites of *MIRIAL*. Employing *TriplexAligner*^36^, we predicted the triplex formation of *MIRIAL* with a selected list of promoters of genes that were regulated following *MIRIAL* knockdown **(Fig. 5B)** . The *FOXO1* promoter region emerged as the highest-scoring candidate in this preliminary, biased analysis. In a more unbiased, genome-wide approach, we identified the top 50 enhancers **(Fig. 5C)** (here denoted as regulatory elements (REMs) and annotated in EpiRegio (https://epiregio.de/)^40, 41^) that are predicted to be target sites of *MIRIAL* triplex formation. These regions were sorted based on their *TriplexAligner* scores (>50) and further stratified based on coverage with active chromatin marks (H3K27ac) in endothelial cells obtained from *ENCODE*^42^. This analysis provides a comprehensive view of the putative *MIRIAL* triplex target-ome in human endothelial cells. Interestingly, the *MIRIAL*-*FOXO1* interaction is in the top 0.15% of all predicted interactions with REMs and promoters combined and when stratified by H3K27ac signal **(Supp. Fig. 6C)** . In addition, the putative *MIRIAL* triplex target site in the *FOXO1* promoter region is not annotated as a transposable element **(Supp. Fig. 6D**, repeats track). However, it (a) is distinct from transcription factor binding sites at the *FOXO1* promoter, (b) encompasses multiple REMs and (c) is an active regulatory region highly enriched in H3K27ac and H3K4me3 histone marks, which are indicative of its likely involvement in the control of *FOXO1* gene expression **(Supp. Fig. 6D)**.

To validate the functionality of *MIRIAL* as a triplex-forming RNA, we employed CRISPR/Cas9 to delete the *MIRIAL* TFR from its genomic locus. As the predicted TFR is located within a stretch of repetitive elements, guide RNAs (gRNAs) were strategically designed for the regions both upstream and downstream of the *Alu* sequence. This targeted knockout strategy resulted in the deletion of 1016 bp **(Fig. 5D)** . We confirmed the knockout in HUVECs by genomic DNA PCR **(Fig. 5E)** and qRT-PCR **(Fig. 5F)**. Our findings indicate that the excision of the transposable elements - including the predicted TFR - from the *MIRIAL* locus results in a downregulation of *FOXO1* **(Fig. 5G)** and an upregulation of *MYC* expression **(Fig. 5H)**, a change comparable to the effects observed following depletion of the full-length *MIRIAL* transcript **(Fig. 2G)** . Furthermore, removal of the TFR triggered an increase of mtDNA content in the cells **(Fig. 5I)**. Our results provide evidence that the four transposable elements within *MIRIAL* harbor a functional region, that appears to influence gene expression and mitochondrial biogenesis, potentially through a direct interaction with chromatin.

### 3.6. *MIRIAL* recruits Mortality Factor 4 Like 2 (MORF4L2) and TATA-binding Protein-associated Factor 2N (TAF15) to the *FOXO1* transcriptional start site

To assess the mechanism underlying the transcriptional activation of *FOXO1* by *MIRIAL*, we used RNA pulldown followed by mass spectrometry to identify protein interaction partners of *MIRIAL*. Our findings validated the interaction of the RNA helicase DDX5/p68 and *MIRIAL* in endothelial cells, which had previously been described in HEK293T cells^12^. Among the enriched proteins, we identified MORF4L2 as a potential interactor of *MIRIAL* **(Fig. 6A)** . MORF4L2 (also known as MRGX) is a component of the NuA4 histone acetyltransferase (HAT) complex^43^. This highly conserved complex is a recognized transcriptional regulator of *FOXO1* in HeLa cells and of the FOXO transcription factor *daf-16* in *C. elegans.* In both species, it has been linked to oxidative stress resistance and contributes to longevity in *C. elegans*^44^. Interestingly, other proteins from this HAT complex (the helicases RUVBL1 and 2, ACTB and ACTL6A/BAF53) were found in the RNA pulldown, although they were not enriched to the same level as MORF4L2.

**Fig. 6:**
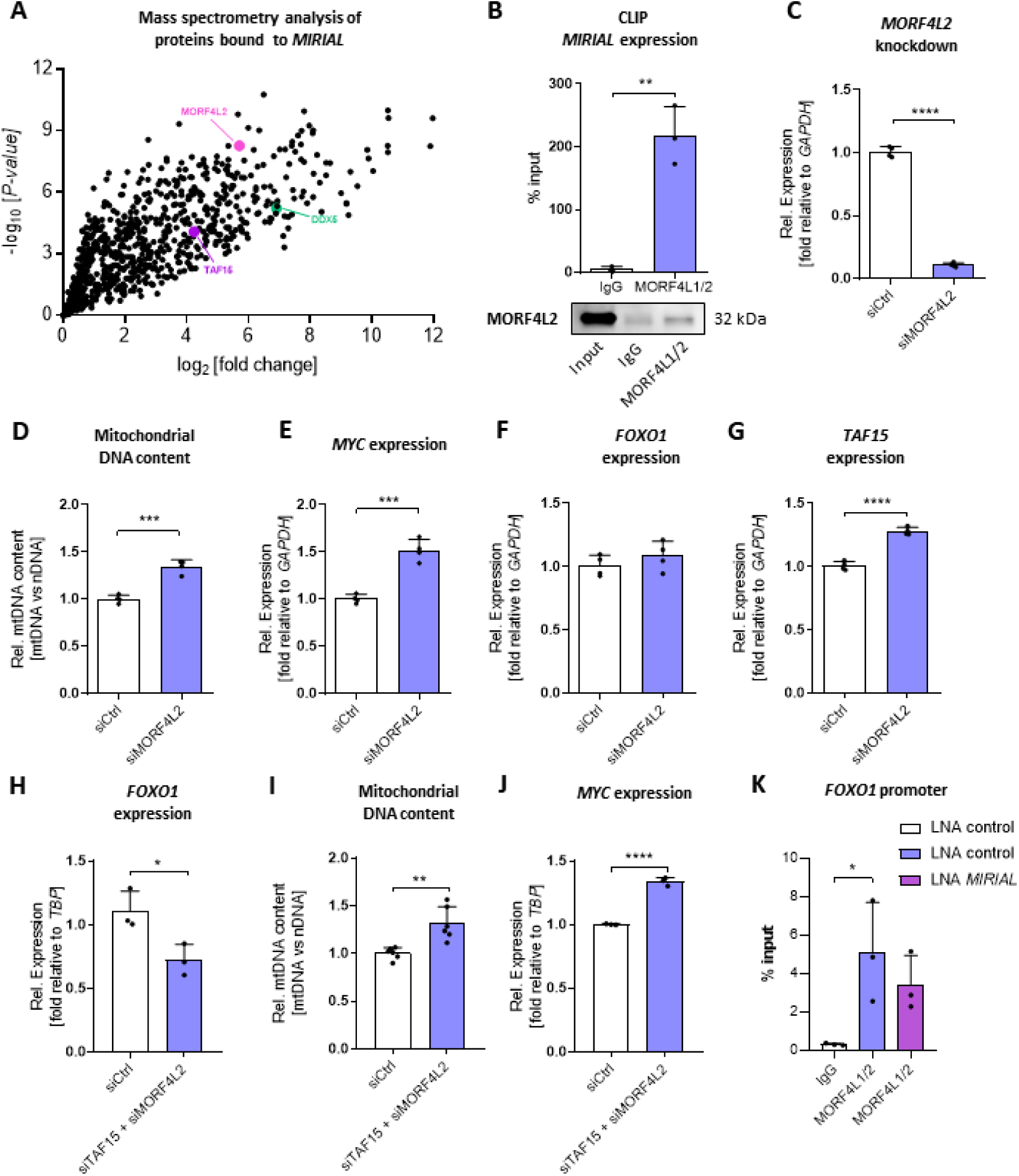
*MIRIAL* recruits Mortality Factor 4 Like 2 (MORF4L2) and TATA-binding Protein-associated Factor 2N (TAF15) to the*FOXO1* transcriptional start site. **A** Volcano plot depicting all proteins from nuclear lysates enriched in *MIRIAL* pulldown as compared to pulldown with a scrambled oligo, identified by mass spectrometry (*n*=6; unpaired *t* test). **B** *MIRIAL* expression measured by qRT-PCR after crosslinking and RNA immunoprecipitation (CLIP) of HUVEC lysate with a MORF4L1/2 antibody or IgG isotype control antibody (5 µg each) and Western blot analysis of MORF4L2 in the input and immunoprecipitates (*n*=3; unpaired *t* test). **C** *MORF4L2* expression in HUVECs 48h after siRNA-mediated *MORF4L2* knockdown (10 nM) compared to control-transfected HUVECs using qRT-PCR (*n*=4; unpaired *t* test). **D** Mitochondrial DNA (mtDNA) content relative to nuclear DNA (nDNA) content in HUVECs 48h after siRNA-mediated *MORF4L2* knockdown (10 nM) compared to control-transfected HUVECs using qPCR (*n*=4; unpaired *t* test). **E** *MY C* expression in HUVECs 48h after siRNA-mediated *MORF4L2* knockdown (10 nM) compared to control-transfected HUVECs using qRT-PCR (*n*=4; unpaired *t* test). **F** *FOX O 1*expression in HUVECs 48h after siRNA-mediated *MORF4L2* knockdown (10 nM) compared to control-transfected HUVECs using qRT-PCR (*n*=4; unpaired *t* test). **G** *TAF15* expression in HUVECs 48h after siRNA-mediated *MORF4L2* knockdown (10 nM) compared to control-transfected HUVECs using qRT-PCR (*n*=4; unpaired *t* test). **H** *FOXO1* expression in HUVECs 48h after siRNA-mediated knockdown of *MORF4L2* and *TAF15* (10 nM each) compared to control-transfected HUVECs using qRT-PCR (*n*=3; unpaired *t* test). **I** Mitochondrial DNA (mtDNA) content relative to nuclear DNA (nDNA) content in HUVECs 48h after siRNA-mediated knockdown of *MORF4L2* and *TAF15* (10 nM each) compared to control-transfected HUVECs using qPCR (*n*=6; unpaired *t* test). **J** *MYC* expression in HUVECs 48h after siRNA-mediated knockdown of *MORF4L2* and *TAF15* (10 nM each) compared to control-transfected HUVECs using qRT-PCR (*n*=3; unpaired *t* test). **K** *FOXO1* promoter sequence measured by qPCR in HUVEC nuclear lysates after chromatin immunoprecipitation (ChIP) using a MORF4L1/2 antibody (5 µg) or IgG isotype control antibody (1 µg). ChIP was performed 48 h after gapmeR-mediated knockdown of *MIRIAL* (20 nM) and compared to control-transfected HUVECs (*n*=3; ANOVA). Results are expressed as mean ± s.d.; \**p*<0.05, \*\**p*<0.01, \*\*\**p*<0.001, \*\*\*\**p*<0.0001

We confirmed the interaction between MORF4L2 and *MIRIAL* by CLIP **(Fig. 6B)** . siRNA-mediated knockdown of *MORF4L2* **(Fig. 6C and Supp. Fig. 7A)** resulted in a significant upregulation of mtDNA content **(Fig. 6D)**, comparable to the previously observed effect of *MIRIAL* silencing **(Fig. 2G)**. Additionally, an upregulation of *MYC* could be observed following MORF4L2 depletion **(Fig. 6E)**, but there was no discernable downregulation of *FOXO1* expression **(Fig. 6F)** . Knockdown of *MORF4L2* was sufficient to upregulate the expression of another potential protein interactor of *MIRIAL*, TAF15 (**Fig. 6A and 6G**). This protein is a component of the transcription initiation factor TF_II_D complex which makes up part of the RNA polymerase II pre-initiation complex^45^. The upregulation of *TAF15* could potentially compensate for the loss of *MORF4L2* . Of note, neither *TAF15* nor *MORF4L2* expression were changed following *MIRIAL* knockdown **(Supp. Fig. 7B** and **7C)** . However, a knockdown of *TAF15* using siRNA **(Supp. Fig. 7D** and **7E)** alongside *MORF4L2* knockdown was sufficient to significantly reduce *FOXO1* expression **(Fig. 6H)** to similar levels as *MIRIAL* knockdown. Moreover, mtDNA content **(Fig. 6I)** and *MYC* expression **(Fig. 6J)** were increased after the double knockdown of *TAF15* and *MORF4L2*, confirming that both proteins are essential for the effects of *MIRIAL* on mitochondria.

To verify that *MIRIAL* recruits MORF4L2 and the NuA4 complex to the *FOXO1* transcriptional start site we performed chromatin immunoprecipitation (ChIP) using a MORF4L1/2 antibody and measured the *FOXO1* promoter sequence in the precipitate by qPCR. We found that MORF4L1/2 interact with the *FOXO1* promoter region **(Fig. 6K)**, and that this interaction was reduced after silencing of *MIRIAL* **(Fig. 6K)** showing that *MIRIAL* helps guiding the NuA4 complex to its genomic target site.

In conclusion, *MIRIAL* interacts with MORF4L2, a component of the NuA4 histone acetyltransferase complex. Through this interaction *MIRIAL* guides this HAT complex to the *FOXO1* promoter region to facilitate *FOXO1* transcription.

### 3.7. *Mirial* KO in mice decreases EF after acute myocardial infarction and regulates angiogenic sprouting *ex vivo*

Having elucidated the molecular mechanism of *MIRIAL* in HUVECs, we questioned whether *Mirial* depletion would have any *in vivo* consequences in a mouse model. To determine its involvement in angiogenesis *in vivo*, we created a *Mirial* knockout (KO) mouse line utilizing CRISPR/Cas9 **(Supp. Fig. 8A)** . We confirmed the successful deletion of *Mirial* in these mice **(Supp. Fig. 8B)**, while the expression of the adjacent miRNA cluster remained unaffected **(Supp. Fig. 8C)** . *Mirial* KO mice exhibited normal viability, fertility and showed no apparent basal defects. Interestingly, in contrast to the *in vitro* findings in HUVECs, we observed that the deletion of *Mirial* did not result in an elevation of mitochondrial content in the hearts of young or aged *Mirial* KO mice compared to their wild-type littermates **(Fig. 7A)** .

**Fig. 7:**
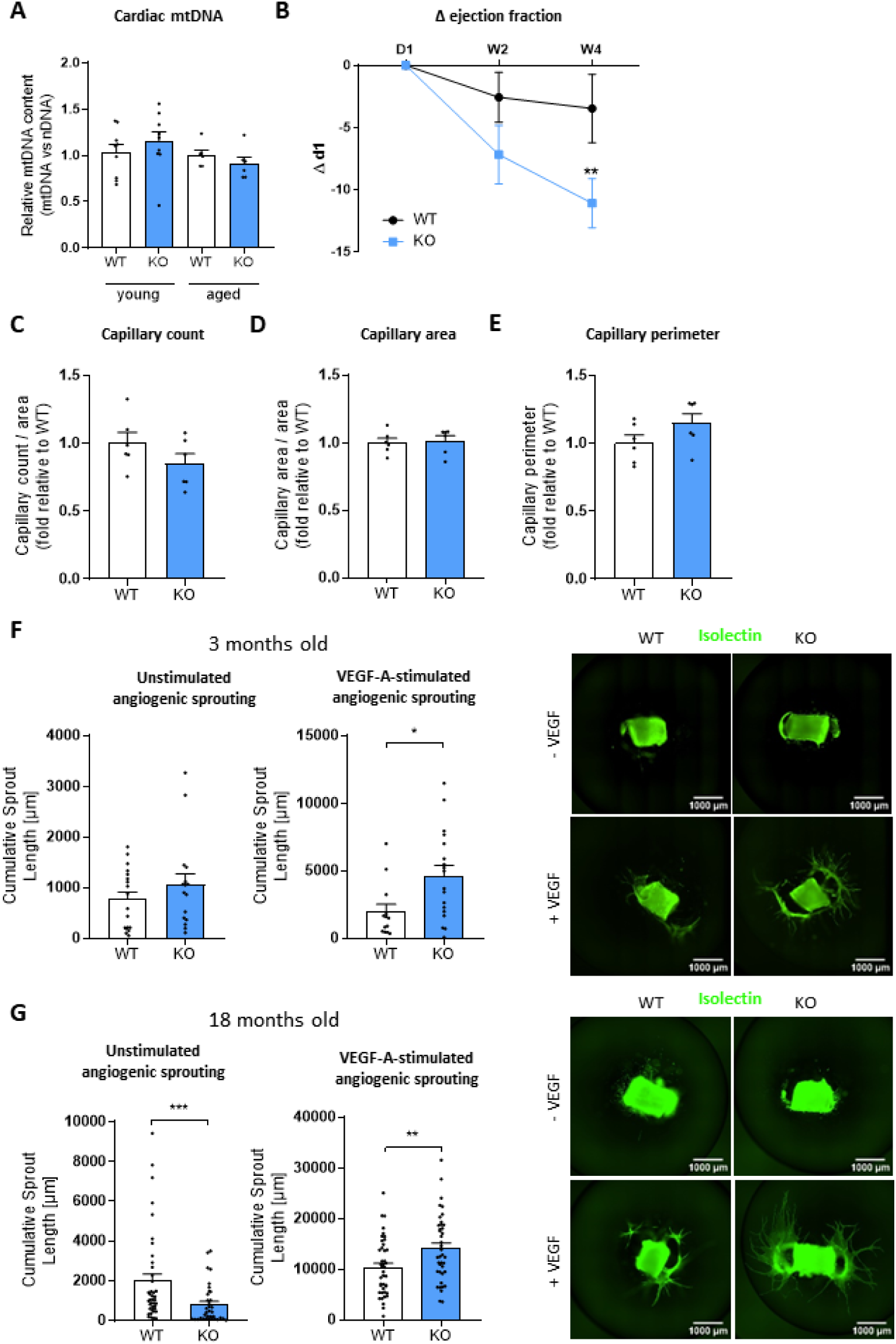
*Mirial* KO in mice decreases EF after acute myocardial infarction and regulates angiogenic sprouting *ex vivo*. **A** Mitochondrial DNA (mtDNA) content relative to nuclear DNA (nDNA) content in heart tissue of young (3 months) and aged (18 months) *Mirial* KO mice and their WT littermates using qPCR (*n*=9 and 6 mice per group; ANOVA). **B** Cardiac contractile function of *Mirial* KO mice and their wildtype (WT) littermates assessed by echocardiography displayed as delta ejection fraction values two weeks (W2) or four weeks (W4) after AMI in comparison to day 1 (D1) (*n*=6; ANOVA). **C** Capillary count in serial cardiac sections of the infarct border zone 4 weeks after AMI surgery of *Mirial* KO mice (11-13 weeks) and their wildtype (WT) littermates. ECs were stained using Isolectin B4 and analyzed using confocal microscopy. Representative images are shown in supp. fig. 8E. Per mouse, three sections were analyzed (*n*=6; unpaired *t* test). **D** Capillary area in serial cardiac sections of the infarct border zone 4 weeks after AMI surgery of *Mirial* KO mice (11-13 weeks) and their wildtype (WT) littermates. ECs were stained using Isolectin B4 and analyzed using confocal microscopy. Representative images are shown in supp. fig. 8E. Per mouse, three sections were analyzed (*n*=6; Mann-Whitney test). **E** Capillary perimeter in serial cardiac sections of the infarct border zone 4 weeks after AMI surgery of *Mirial* KO mice (11-13 weeks) and their wildtype (WT) littermates. ECs were stained using Isolectin B4 and analyzed using confocal microscopy. Representative images are shown in supp. fig. 8E. Per mouse, three sections were analyzed (*n*=6; unpaired *t* test). **F** Cumulative sprout length from aortic rings of young (3 months) *Mirial* KO mice and WT littermates under basal and VEGF-stimulated (30 ng/mL) conditions (*n*=6 mice per group; Mann-Whitney test). ECs were stained with Isolectin B4 and analyzed using immunofluorescence microscopy. 2-8 aortic rings per mouse were analyzed. **G** Cumulative sprout length from aortic rings of aged (18 months) *Mirial* KO mice and WT littermates under basal and VEGF-stimulated (30 ng/mL) conditions (*n*=6 mice per group; Mann-Whitney test and unpaired *t* test). ECs were stained with Isolectin B4 and analyzed using immunofluorescence microscopy. 5-8 aortic rings per mouse were analyzed. Results are expressed as mean ± s.e.m.; \**p*<0.05, \*\**p*<0.01, \*\*\**p*<0.001

To evaluate the impact of *Mirial* KO on the restoration of myocardial capillarization and perfusion post-injury, we conducted acute myocardial infarction (AMI) surgery, followed by echocardiographic assessment of cardiac function and histological examination of the heart. Echocardiography was performed on the first day post-surgery, as well as at two- and four-weeks post-surgery **(Supp. Fig. 8D)**. Comparing the change of contractile function (ΔEF) from the first day post-surgery to the four-week endpoint revealed a significantly reduced ejection fraction in *Mirial* knockout animals **(Fig. 7B)** .

To assess the effects of *Mirial* loss on the cardiac phenotype following AMI at the cellular level, we performed immunohistochemical staining of serial sections of the heart **(Supp. Fig. 8E)**. Capillary density analysis revealed no change in number, size, or perimeter **(Fig. 7C-7E)** of capillaries in the border zone of AMI heart sections. Moreover, histological examination following Sirius Red staining of collagen showed no significant alterations in infarct size or fibrosis after *Mirial* knockout **(Supp. Fig. 8F-8H)**.

Next, *ex vivo* angiogenic capacity was tested using the aortic ring assay with aortic samples from both young (3 months) and aged (18 months) **(Fig. 7F and 7G)** mice. In old animals basal sprouting was drastically reduced after *Mirial* KO. Additionally, confirming our previous observations of the *in vitro* sprouting assay, these *ex vivo* experiments show that endothelial cells from *Mirial* KO animals exhibit increased responsiveness to VEGF-A in in both age brackets.

Together, these experiments show that loss of *Mirial in vivo* has an adverse effect on cardiac outcome after AMI. Moreover, *Mirial* is an important regulator of VEGF-A-response and angiogenic sprouting *in vitro* and *ex vivo*.

## 4. Discussion

The present study reveals that the aging-induced lncRNA *MIRIAL* promotes endothelial cell proliferation and migration, while reducing mitochondrial mass, spare respiratory capacity and p53 signaling through modulation of the FOXO1-MYC antagonism, desensitizing endothelial cells against VEGF-A signaling.

In the present study, *MIRIAL* knockdown in HUVECs and *Mirial* knockout in mice had divergent effects on angiogenic sprouting if VEGF-A was administered compared to basal conditions. Under unstimulated conditions loss of *MIRIAL* decreases sprouting. This phenotype can be attributed to the accompanying induction of the p53 pathway and reduction in migration and proliferation. However, we show that VEGF-A administration not only rescues the effect of *MIRIAL* knockdown on proliferation, but also leads to overshooting angiogenic sprouting, both *in vitro* and *ex vivo*. We hypothesize that this is due to a strong induction of MYC through VEGFR2 signaling^46^, which overrules the effects of p53 signaling.

While there was no change in capillarization, infarct size or fibrosis after AMI, a noticeable reduction in the change of ejection fraction was observed. This may be due to an as-yet-undefined function of *Mirial* in cardiomyocytes, which could potentially enhance or protect contractile function after AMI.

In mice, decreased expression of *Mirial* has been linked to reduced FoxO signaling^8^. In our *Mirial* KO mouse model, the regulation of FoxO signaling could not be confirmed. Comparing the human and mouse orthologues of *MIRIAL* shows that the mouse transcript is significantly shorter than the human orthologue (690 bp versus 1774 bp) and lacks the *Alu* element-containing region, as *Alu* elements are only conserved in primates^47^. We conclude that human *MIRIAL* and mouse *Mirial* are only partially functionally conserved, and that human *MIRIAL* alters gene expression *via* a triplex-dependent mechanism that is not evident in mice.

The putative TFR of *MIRIAL*, that is predicted to interact with the *FOXO1* promoter region, contains an *Alu* element. Interestingly, lncRNAs, like S*MANTIS*^37, 38^ and *ANRIL*^39^, were described to contain *Alu* elements as functional regions and were shown to regulate endothelial cell function and atherogenesis. The lncRNA *KCNQ1OT1* contains a transposable element of the L1 class capable of forming a triplex with L1 and *Alu* elements on genomic DNA^48^. However, unlike the aforementioned examples, our study is the first to describe a triplex-forming *Alu* element within a lncRNA. This unique feature sets *MIRIAL* apart from other lncRNAs that have been studied until now.

## Conclusion

Taken together, *MIRIAL* is an aging-induced lncRNA which acts as a key regulator of endothelial metabolic and cellular function. *MIRIAL* promotes cell proliferation, migration and basal angiogenic sprouting while decreasing mitochondrial function. However, silencing *MIRIAL* in pro-angiogenic conditions improves angiogenesis. We hypothesize that *MIRIAL* influences these cellular functions by affecting the p53 pathway and mitochondrial respiration through FOXO1 signaling. These results suggest that transcriptional modulation of *MIRIAL*, particularly in the elderly, might be a promising strategy to improve therapeutic angiogenesis.

## Acknowledgments and sources of funding

This work was supported by the Goethe University Frankfurt/Main, Germany; the Deutsche Forschungsgemeinschaft (DFG) excellence cluster EXS2026 (Cardio-Pulmonary Institute) [project number 390649896]; the Deutsche Forschungsgemeinschaft (DFG) Transregio project TRR267 [project number 403584255, project B04 to RAB and LM, project B03 and A01 to SD, project A04 to MSL, project A06 to RPB, and project Z02 to IW]; the Deutsche Forschungsgemeinschaft (DFG) collaborative research centre SFB 834 [project number 75732319, project B09 to RAB); the German Centre for Cardiovascular Research (DZHK); an European Research Council (ERC) starting grant (NOVA) [project number 638178]; and an European Research Council (ERC) consolidator grant (NICCA) [project number 101002599]. The graphical abstract was created with BioRender.com.

## Author contributions

CK, KT, and RAB conceptualized and designed the study. CK, SK, TW, KT, AF, RPJ, MMR, DB, FL, FV, JUGW, JS, ATG, LS, SG, IW, LM and MSL acquired, analyzed, and interpreted data. CK drafted the manuscript. All authors critically revised the manuscript and approved the final version to be published.

## Conflict of Interest

non declared.

## List of Supplementary Data

1. Supplementary Figures and Legends 1-8
2. Supplementary Methods
3. Supplementary Tables 1-5
4. Supplementary References

## Notes

### Competing Interest Statement

The authors have declared no competing interest.

### Summary of Updates

We shortened the results and discussion section to improve readability.

