## Supplemental data for "The aging-induced long non-coding RNA *MIRIAL* controls endothelial cell and mitochondrial function"

Kohnle et al.

### 1. Supplementary figures and legends

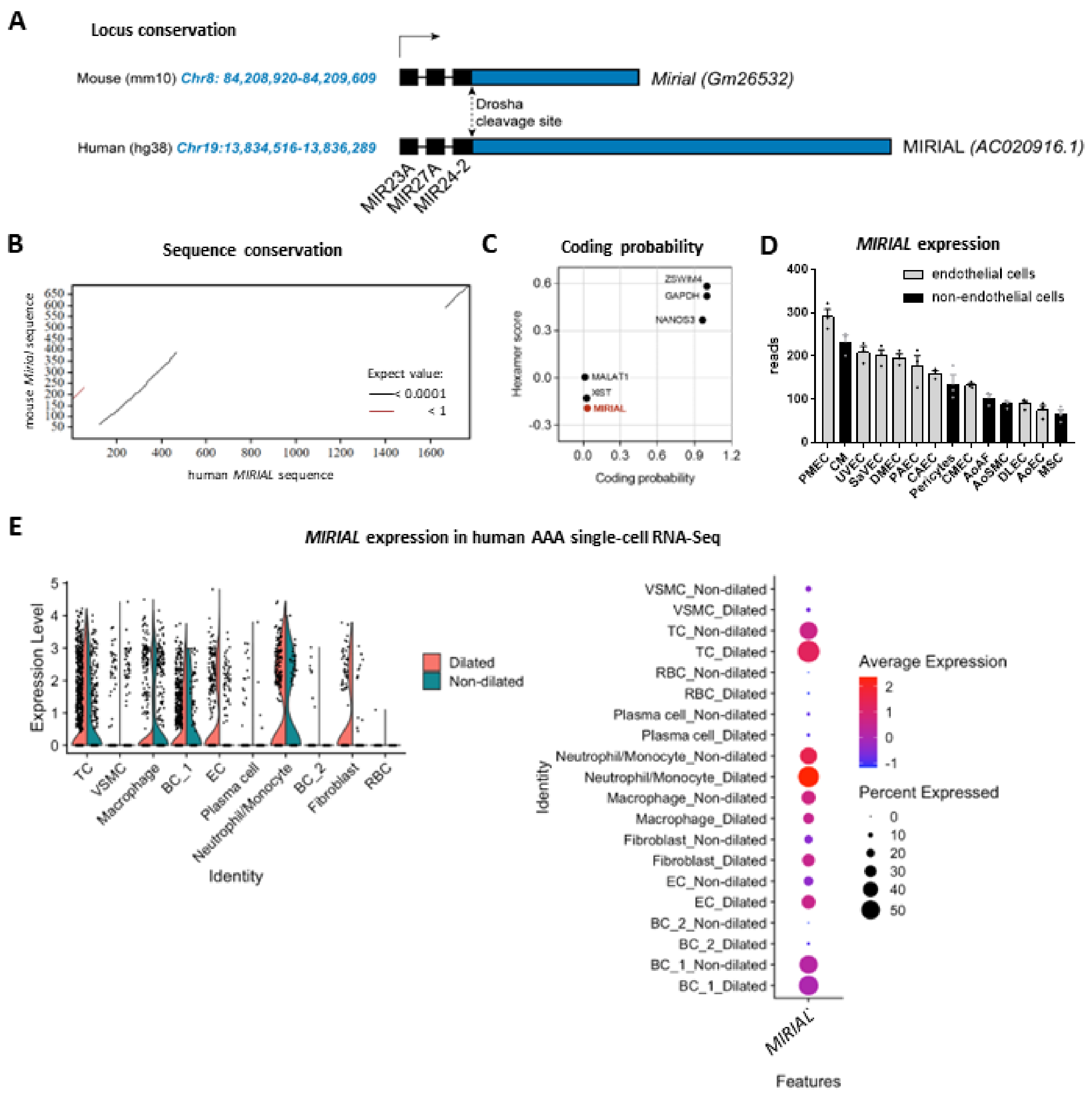


**Supp. Fig. 1: A** Schematic representation of the orthologous *Mirial* (Gm26532) and *MIRIAL* (AC020916.1) loci in mouse and human, respectively, including the chromosome coordinates and the miRNA cluster miR-23A~27A~24-2. **B** Sequence conservation between mouse *Mirial* (y-axis) and human *MIRIAL* (x-axis) using the LALIGN DNA:DNA tool of the FASTA Sequence Comparison software of the University of Virginia. **C** Coding probability of *MIRIAL* compared to established non-coding transcripts (*MALAT1* and *XIST*) and protein-coding transcripts (*GAPDH* and *NANOS3*) as well as the nearest protein-coding gene to the *MIRIAL* locus

(*ZSWIM4*) using the Coding-Potential Assessment Tool (CPAT). **D** *MIRIAL* expression in human endothelial cells (ECs) from different vascular beds (PMEC = Pulmonary Microvascular ECs, UVEC = Umibilical Vein ECs, SaVEC = Saphenous Vein ECs, DMEC = Dermal Microvascular ECs, PAEC = Pulmonary Arterial ECs, HCAEC = Coronary Artery ECs, CMEC = Cardiac Microvascular ECs, DLEC = Dermal Lymphatic ECs, AoEC = Aortic ECs) and non-endothelial cardiovascular cells (CM = cardiomyocytes, pericytes, AoAF = Aortic Arterial Fibroblasts, AoSMC = Aortic Smooth Muscle Cells, MSC = mesenchymal stem cells) measured by RNA sequencing (*n*=3). Results are expressed as mean ± s.e.m. **E** Violin plot and dot plot showing the expression pattern of *MIRIAL* in identified cell clusters in abdominal aortic aneurysm (AAA) patient samples (dilated) compared to undilated control samples from the same patients using single-cell RNA sequencing (TC = T Cells, VSMC = Vascular Smooth Muscle Cells, BC = B cells, EC = Endothelial Cells, RBC = Red Blood Cells) (*n*=4).

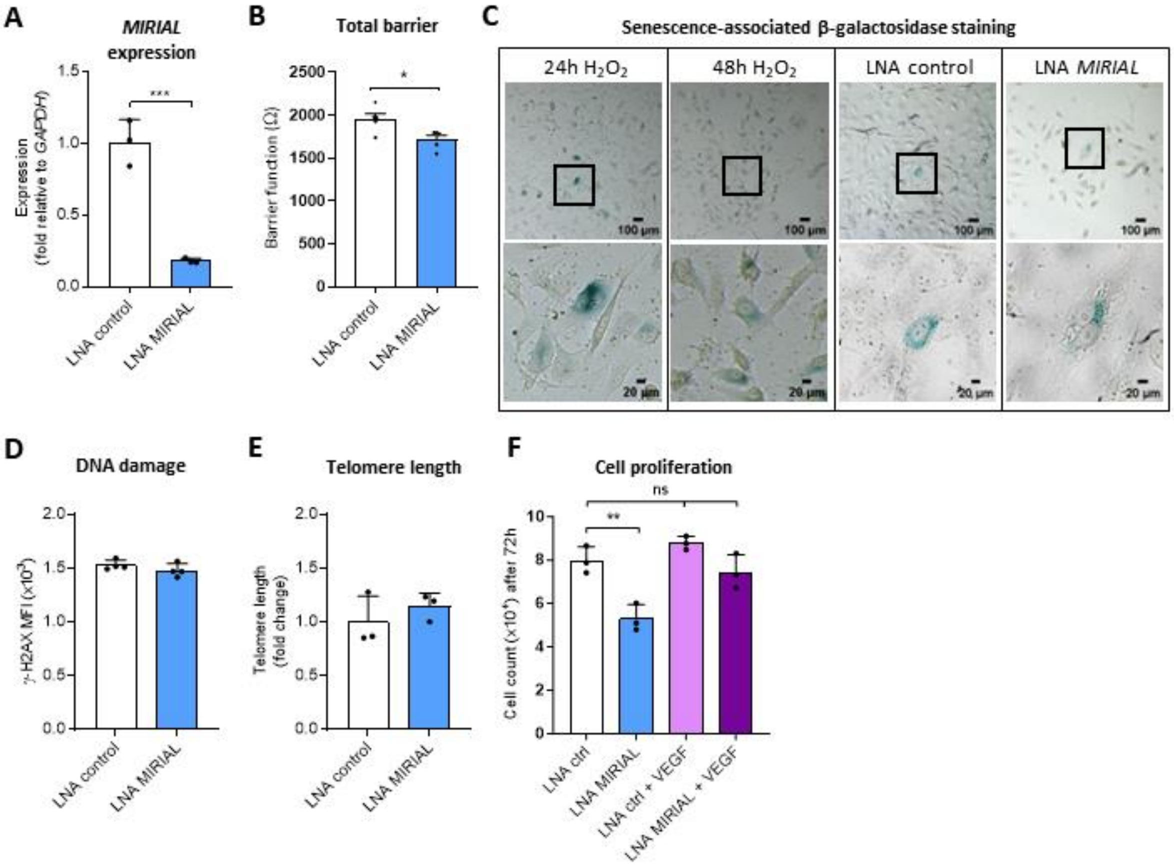


**Supp. Fig. 2: A** *MIRIAL* expression in HUVECs 48h after gapmeR-mediated *MIRIAL* knockdown (20 nM) compared to control-transfected HUVECs using qRT-PCR (*n*=3; unpaired *t* test). **B** Electric cell-substrate impedance sensing (ECIS)-based measurement of the endothelial barrier function of HUVECs 48h after gapmeR-mediated *MIRIAL* knockdown (20 nM) compared to control-transfected HUVECs (*n*=5; unpaired *t* test). **C** Representative images of senescence-associated β-galactosidase staining in HUVECs 48h after gapmeR-mediated *MIRIAL* knockdown (20 nM) compared to control-transfected HUVECs. Treatment of untransfected HUVECs with $H_2O_2$ (200 µM) for 24h or 48h, respectively, was included as a positive control. **D** DNA damage in HUVECs 48h after gapmeR-mediated *MIRIAL* knockdown (20 nM) compared to control-transfected HUVECs measured by γH2A.X (Ser139) staining followed by flow cytometry (*n*=4; unpaired *t* test). **E** Telomere length in HUVECs 48h after gapmeR-mediated *MIRIAL* knockdown (20 nM) compared to control-transfected HUVECs measured by qPCR ($n$=3; unpaired $t$ test). **F** Proliferation of HUVECs measured as cell count 72h post-transfection. HUVECs were transfected with a gapmeR targeting *MIRIAL* or a negative control gapmeR (20 nM) and treated with VEGF-A (50 ng/mL) or vehicle (PBS) ($n$=3; ANOVA). Results are expressed as mean ± s.d.; *$p<0.05$, **$p<0.01$

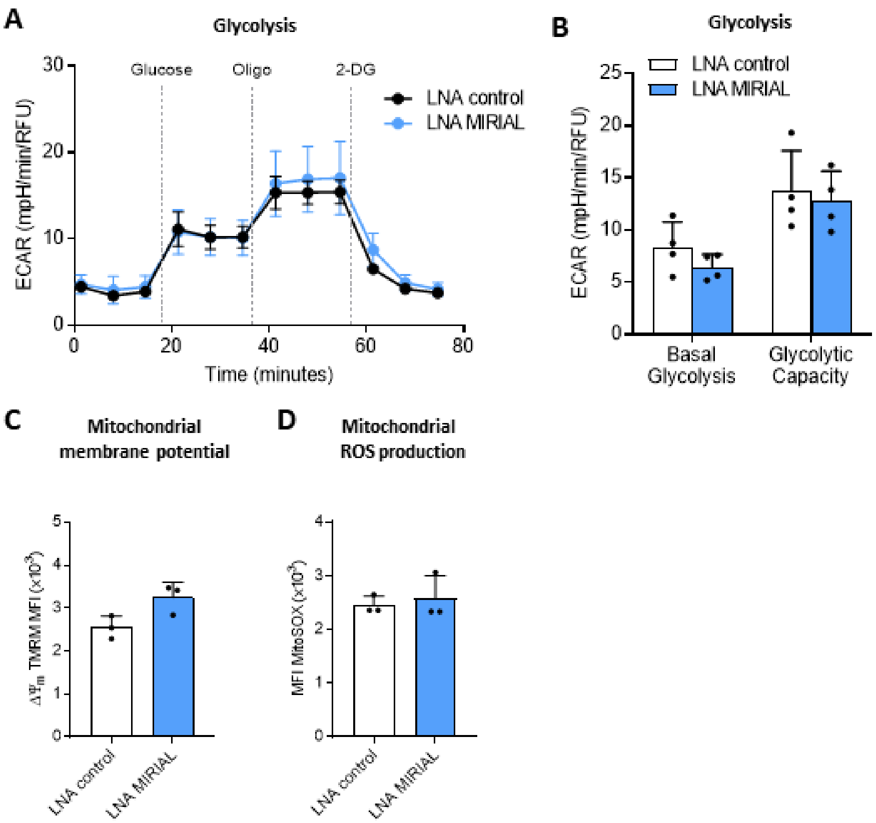


**Supp. Fig. 3: A** Extracellular acidification rate (ECAR) as a measure of glycolysis in HUVECs 48h after gapmeR-mediated *MIRIAL* knockdown (20 nM) compared to control-transfected HUVECs using the Seahorse glycolysis stress test (representative graph of *n*=4). **B** Extracellular acidification rate (ECAR) as a measure of basal glycolysis and glycolytic capacity in HUVECs 48h after gapmeR-mediated *MIRIAL* knockdown (20 nM) compared to control-transfected HUVECs using the Seahorse glycolysis stress test, quantification from supp. fig. 3A (*n*=4; ANOVA). **C** Mitochondrial membrane potential in HUVECs 48h after gapmeR-mediated *MIRIAL* knockdown (20 nM) compared to control-transfected HUVECs measured by tetramethylrhodamine-methylester (TMRM) staining followed by flow cytometry (*n*=3; unpaired *t* test). **D** Mitochondrial production of reactive oxygen species (ROS) in HUVECs 48h after gapmeR-mediated *MIRIAL* knockdown (20 nM) compared to control-transfected HUVECs measured by MitoSOX staining followed by flow cytometry (*n*=3; Mann-Whitney test). Results are expressed as mean ± s.d.

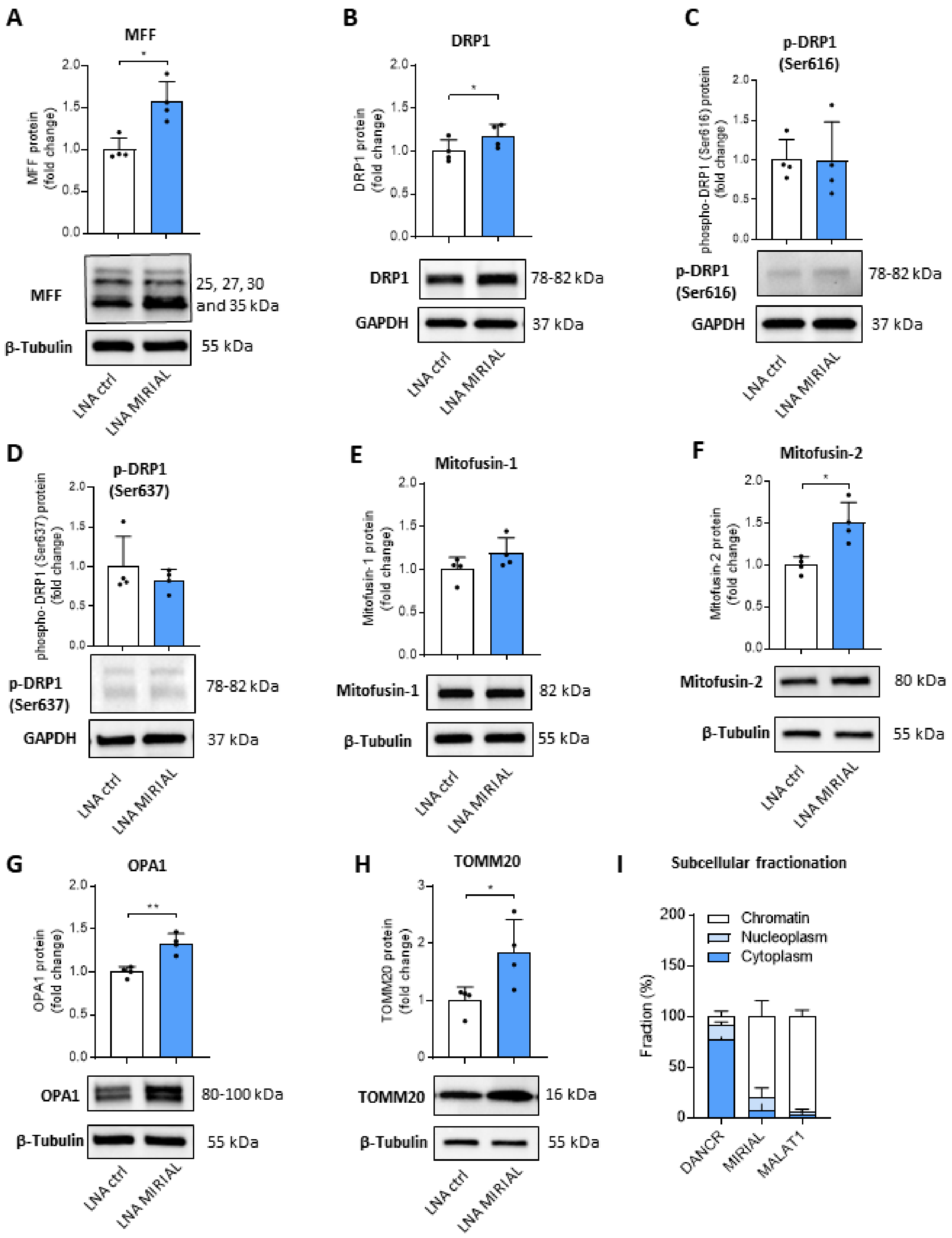


**Supp. Fig. 4: A** Protein levels of MFF in HUVECs 48h after gapmeR-mediated *MIRIAL* knockdown (20 nM) compared to control-transfected HUVECs using Western Blot (*n*=4; Mann-Whitney test). **B** Protein levels of DRP1 in HUVECs 48h after gapmeR-mediated *MIRIAL* knockdown (20 nM) compared to control-transfected HUVECs using Western Blot (*n*=4; paired *t* test). **C** Protein levels of phosphorylated DRP1 (Ser616) in HUVECs 48h after gapmeR-mediated *MIRIAL* knockdown (20 nM) compared to control-transfected HUVECs using Western Blot (*n*=4; paired *t* test). **D** Protein levels of phosphorylated DRP1 (Ser637) in HUVECs 48h after gapmeR-mediated *MIRIAL* knockdown (20 nM) compared to control-transfected HUVECs using Western Blot (*n*=4; Mann-Whitney test). **E** Protein levels of Mitofusin-1 in HUVECs 48h after gapmeR-mediated *MIRIAL* knockdown (20 nM) compared to control-transfected HUVECs using Western Blot (*n*=4; paired *t* test). **F** Protein levels of Mitofusin-2 in HUVECs 48h after gapmeR-mediated *MIRIAL* knockdown (20 nM) compared to control-transfected HUVECs using Western Blot (*n*=4; paired *t* test). **G** Protein levels of OPA1 in HUVECs 48h after gapmeR-mediated *MIRIAL* knockdown (20 nM) compared to control-transfected HUVECs using Western Blot (*n*=4; paired *t* test). **H** Protein levels of TOMM20 in HUVECs 48h after gapmeR-mediated *MIRIAL* knockdown (20 nM) compared to control-transfected HUVECs using Western Blot (*n*=4; Mann-Whitney test). **I** Expression of the lncRNAs *DANCR*, *MIRIAL* and *MALAT1* in HUVEC subcellular fractions (cytoplasm, nucleoplasm and chromatin) using qRT-PCR (*n*=3). Results are expressed as mean ± s.d.; *$p<0.05$, **$p<0.01$

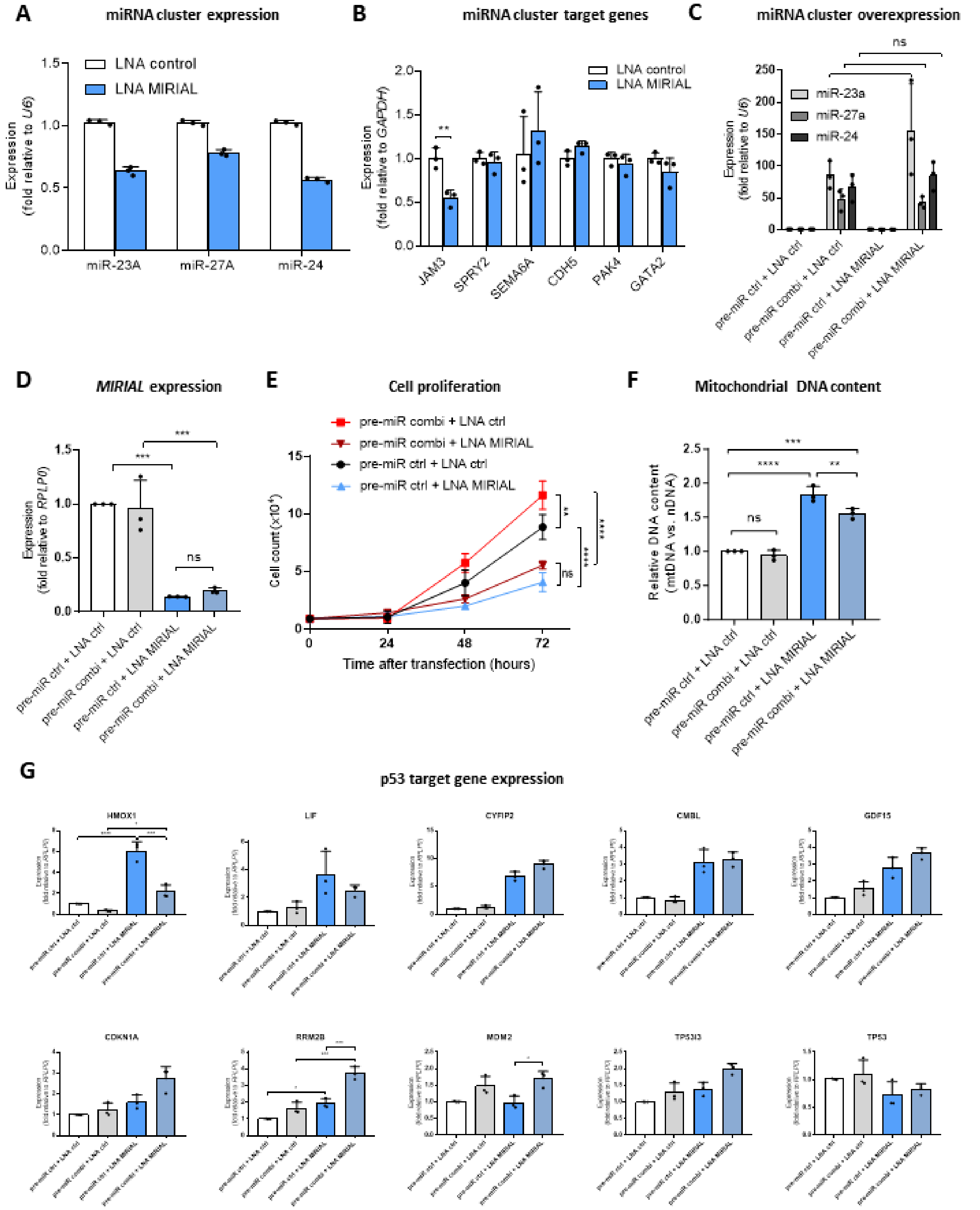


**Supp. Fig. 5: A** Expression of the miRNAs miR-23A, miR-27A and miR-24-2 in HUVECs 48h after gapmeR-mediated *MIRIAL* knockdown (20 nM) compared to control-transfected HUVECs measured using TaqMan qRT-PCR ($n$=3; Kruskal-Wallis test). **B** Expression of selected target genes of miR-23A (*JAM3* and *SPRY2*), miR-27A (*SEMA6A* and *CDH5*) and miR-24-2 (*PAK4* and

*GATA2*) in HUVECs 48h after gapmeR-mediated *MIRIAL* knockdown (20 nM) compared to control-transfected HUVECs measured using qRT-PCR ($n$=3; ANOVA). **C** Overexpression of the miRNAs miR-23A, miR-27A and miR-24-2 measured using TaqMan qRT-PCR. HUVECs were transfected for 48h with a mix of equal amounts (10 nM each) of precursor miRNAs of miR-23A, miR-27A and miR-24-2 (pre-miR combi) or a negative pre-miR-control (30 nM) and a gapmeR targeting *MIRIAL* or a negative control gapmeR (20 nM) ($n$=3; ANOVA). **D** *MIRIAL* expression using qRT-PCR. HUVECs were transfected for 48h with a mix of equal amounts (10 nM each) of precursor miRNAs of miR-23A, miR-27A and miR-24-2 (pre-miR combi) or a negative pre-miR-control (30 nM) and a gapmeR targeting *MIRIAL* or a negative control gapmeR (20 nM) ($n$=3; ANOVA). **E** Growth curve depicting the cell number of HUVECs after the indicated hours (0-72h) post-transfection. HUVECs were transfected with a mix of equal amounts (10 nM each) of precursor miRNAs of miR-23A, miR-27A and miR-24-2 (pre-miR combi) or a negative pre-miR-control (30 nM) and a gapmeR targeting *MIRIAL* or a negative control gapmeR (20 nM) (representative graph of $n$=3; ANOVA). **F** Mitochondrial DNA (mtDNA) content relative to nuclear DNA (nDNA) content in HUVECs 48h post-transfection using qPCR. HUVECs were transfected with a mix of equal amounts (10 nM each) of precursor miRNAs of miR-23A, miR-27A and miR-24-2 (pre-miR combi) or a negative pre-miR-control (30 nM) and a gapmeR targeting *MIRIAL* or a negative control gapmeR (20 nM) ($n$=3; ANOVA). **G** Expression of *TP53* and p53 target genes regulated after *MIRIAL* knockdown (fig. 3A and 3C) in HUVECs 48h post-transfection using qRT-PCR. HUVECs were transfected for 48h with a mix of equal amounts (10 nM each) of precursor miRNAs of miR-23A, miR-27A and miR-24-2 (pre-miR combi) or a negative pre-miR-control (30 nM) and a gapmeR targeting *MIRIAL* or a negative control gapmeR (20 nM) ($n$=3; ANOVA or Kruskal-Wallis test). Results are expressed as mean ± s.d.; *$p$<0.05, **$p$<0.01, ***$p$<0.001, ****$p$<0.0001

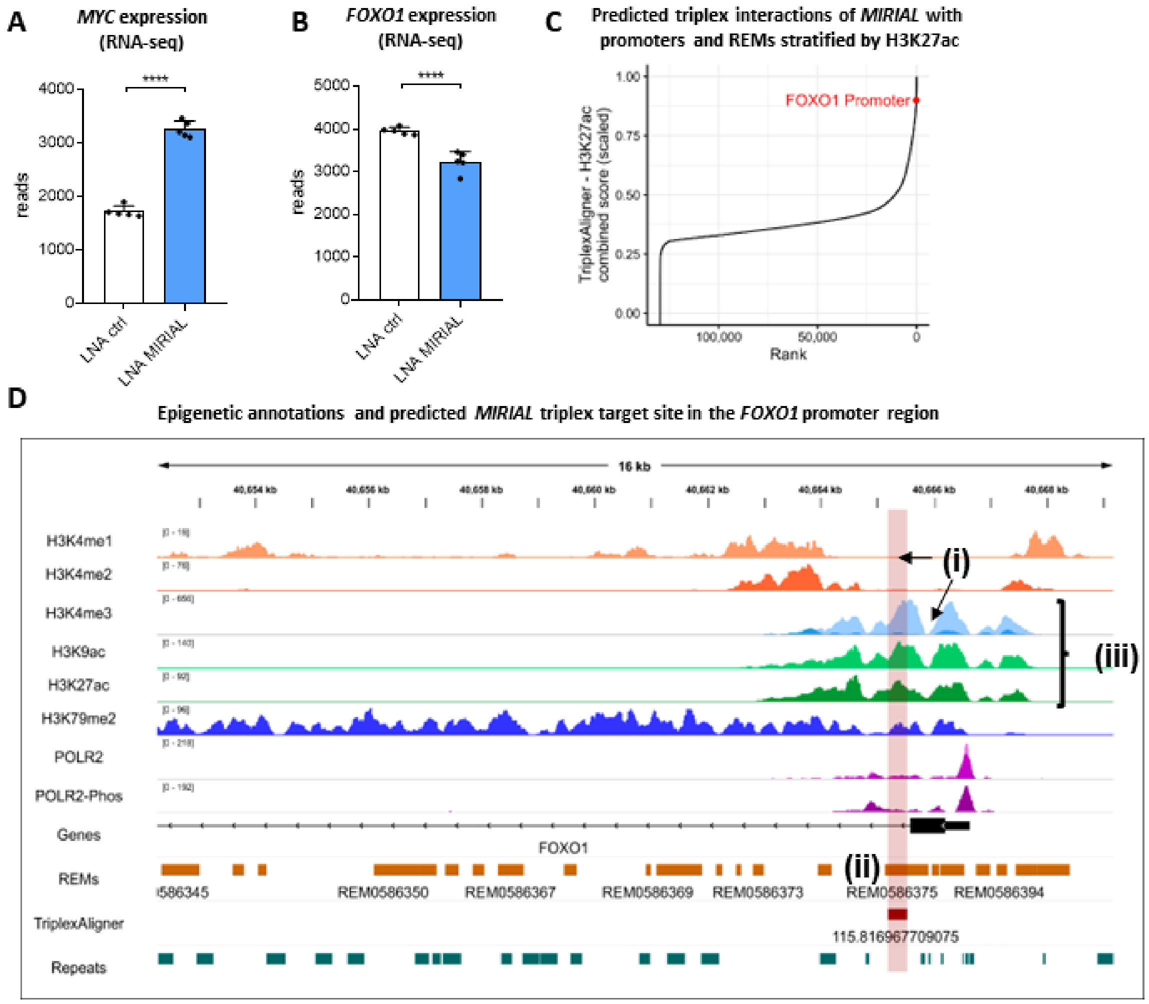


**Supp. Fig. 6: A** *MYC* expression in HUVECs 48h after gapmeR-mediated *MIRIAL* knockdown (20 nM) compared to control-transfected HUVECs measured by RNA sequencing (*n*=6; Mann-Whitney test with FDR correction). **B** *FOXO1* expression in HUVECs 48h after gapmeR-mediated *MIRIAL* knockdown (20 nM) compared to control-transfected HUVECs measured by RNA sequencing (*n*=6; Mann-Whitney test with FDR correction). **C** All predicted interactions of *MIRIAL* with regulatory elements (REMs) and promoters, stratified by H3K27ac signal. When *TriplexAligner* score and H3K27ac signal in the triplex site are combined, the *FOXO1* promoter region target site of *MIRIAL* is in ranked #188 (top 0.15%) of more than 150,000 potential *MIRIAL* target regions. Scaled score = $\log_{10}$(*TriplexAligner* $-\log_{10}$(E) × H3K27ac signal in target site) scaled to between 0 & 1. **D** Epigenetic annotations and predicted *MIRIAL* triplex target site in the *FOXO1* promoter region illustrated using the Integrative Genomics Viewer

(IGV). (i) The *MIRIAL* target site is distinct from the transcription factor binding sites at the *FOXO1* promoter. (ii) The predicted *MIRIAL* target site in the *FOXO1* promoter region covers multiple REMs. (iii) The target site is an active regulatory region with chromatin highly decorated with H3K27ac, H3K9ac and H3K4me3. Results are expressed as mean ± s.d.; ****$p<0.0001$

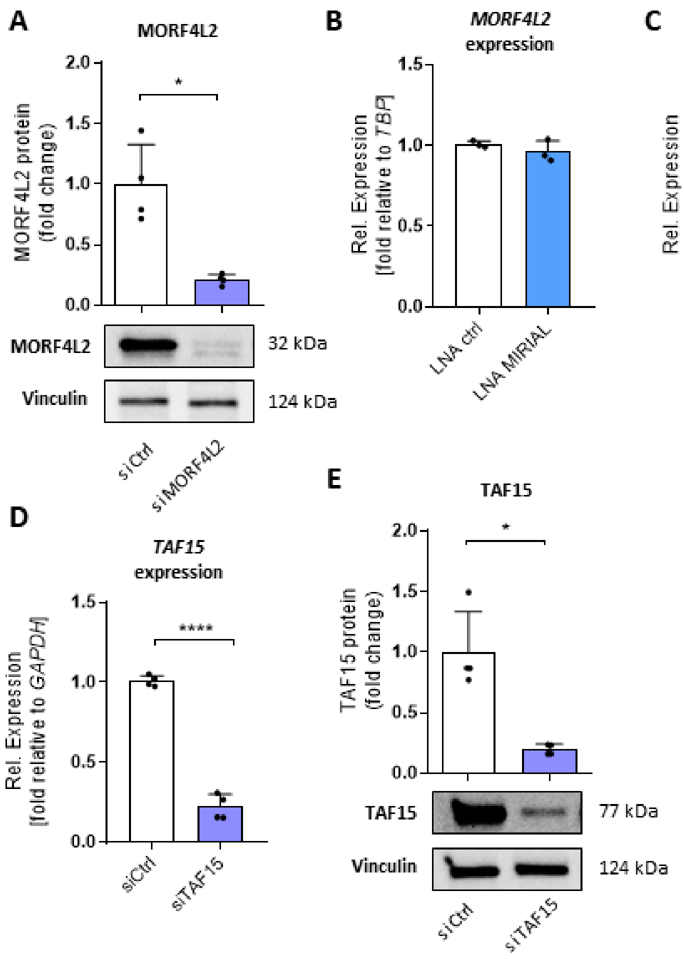


**Supp. Fig. 7: A** Protein levels of MORF4L2 in HUVECs 48h after siRNA-mediated *MORF4L2* knockdown (10 nM) compared to control-transfected HUVECs using Western Blot (*n*=4; paired *t* test). **B** *MORF4L2* expression in HUVECs 48h after gapmeR-mediated *MIRIAL* knockdown (20 nM) compared to control-transfected HUVECs using qRT-PCR (*n*=3; unpaired *t* test). **C** *TAF15* expression in HUVECs 48h after gapmeR-mediated *MIRIAL* knockdown (20 nM) compared to control-transfected HUVECs using qRT-PCR (*n*=3; unpaired *t* test). **D** *TAF15* expression in HUVECs 48h after siRNA-mediated *TAF15* knockdown (10 nM) compared to control-transfected HUVECs using qRT-PCR (*n*=4; unpaired *t* test). **E** Protein levels of TAF15 in HUVECs 48h after siRNA-mediated *TAF15* knockdown (10 nM) compared to control-transfected HUVECs using Western Blot (*n*=4; Mann-Whitney test). Results are expressed as mean ± s.d.; *$p$<0.05, ****$p$<0.0001

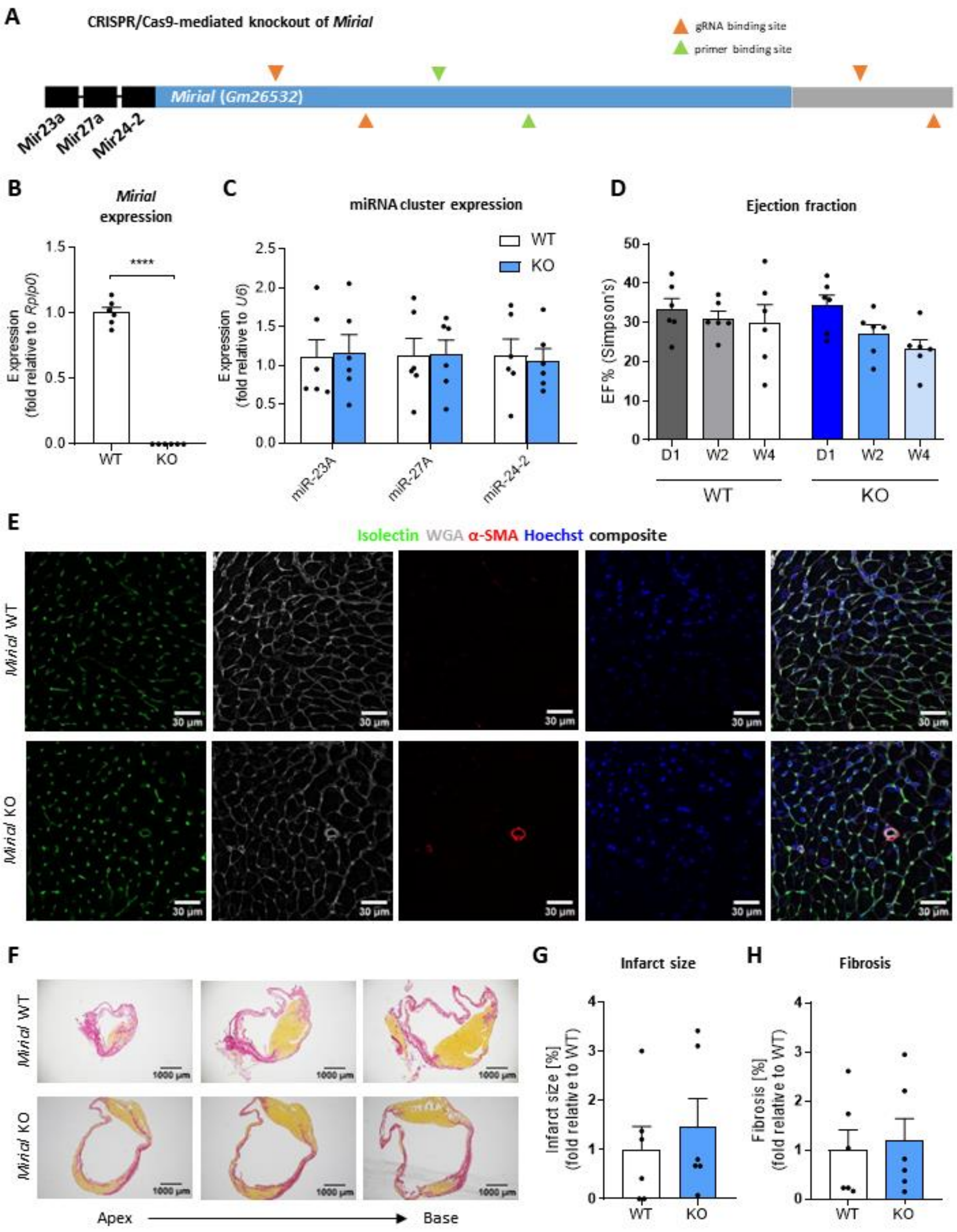


**Supp. Fig. 8: A** Schematic representation of the *Mirial* locus including the miR-23a~27a~24-2 cluster and the binding sites of guide RNAs (gRNAs) used for CRISPR/Cas9-mediated knockout (KO) of *Mirial* in mice as well as primer binding sites to detect the KO using qRT-PCR. **B** *Mirial* expression in the lungs of *Mirial* KO mice (13-14 weeks) and their wildtype (WT) littermates using qRT-PCR ($n=6$; unpaired $t$ test). **C** Expression of the miRNAs miR-23A, miR-27A and miR-24-2 in the lungs of *Mirial* KO mice (13-14 weeks) and their wildtype (WT) littermates using TaqMan qRT-PCR ($n=6$; ANOVA). **D** Cardiac contractile function of *Mirial* KO mice and their wildtype (WT) littermates assessed by echocardiography on day one (D1), two weeks (W2) or four weeks (W4) after AMI surgery displayed as percent ejection fraction values ($n=6$; unpaired $t$ test). **E** Representative images of serial sections of the heart of *Mirial* KO mice and their WT littermates four weeks after AMI surgery stained with Isolectin (ECs), wheat germ agglutinin (WGA) (all cell membranes), anti-alpha smooth muscle actin (SMA) (vascular smooth muscle cells of larger vessels) and Hoechst33342 (nuclei). Images were obtained from the infarct border zone using confocal microscopy. **F** Representative images of serial sections of the heart of *Mirial* KO mice and their WT littermates four weeks after AMI surgery stained with Sirius Red. **G** Infarct size quantified from serial sections of the heart of *Mirial* KO mice and their WT littermates four weeks after AMI surgery stained with Sirius Red and shown in supp. fig. 8F. Infarct size was assessed by measuring the length of the infarcted region as percentage of the left ventricular circumference ($n=6$; unpaired $t$ test). **H** Fibrosis quantified from serial sections of the heart of *Mirial* KO mice and their WT littermates four weeks after AMI surgery stained with Sirius Red and shown in supp. fig. 8F. Fibrosis was assessed by measuring the positively stained area as percentage of the total area of the section ($n=6$; unpaired $t$ test). Results are expressed as mean ± s.e.m.; ****$p<0.0001$

### 2. Supplementary Methods

#### Aortic ring assay

Aortic rings were prepared as described before [2]. Aortic rings were fed with fresh growth medium containing VEGF-A (PeproTech) every 2-3 days for 7 days. Then the rings were fixed with Histofix (Roth), blocked with Protein Block Serum-Free (Dako) and subsequently stained with Isolectin B4 (Vector) and Streptavidin Alexa488 (Invitrogen) and imaged using the Nikon Eclipse Ti2 microscope with 10x magnification and in 5x5 large image mode. Cumulative sprout length was determined using Image J software.

#### ATP determination assay

Cellular ATP levels were assessed utilizing the Molecular Probes ATP determination kit (Invitrogen). The assay was carried out 48 h after transfection of HUVECs and in accordance with the manufacturer's guidelines. After washing with PBS (Gibco), cells were detached, counted, washed, and resuspended at a concentration of 1,000 cells/µL. Cells were boiled at 95°C for 10 min to achieve cell lysis. Subsequently, the samples were transferred into a white-walled 96-well plate (Greiner) and mixed with the reaction solution supplied with the kit. Luminescence was measured in the samples and an ATP standard curve using the Glomax Multi+ Detection System multiplate reader (Promega).

#### Caspase assay

24 h after transfection, cells were re-seeded into black-walled 96-well plates (Corning) and caspase activity was determined additional 24 h later using the Apo-ONE Homogeneous Caspase-3/7 kit (Promega) according to the manufacturer’s instructions. Fluorescence was

measured using the Glomax Multi+ Detection System plate reader by Promega (excitation 485+/- 20 nm, emission 530 +/- 25 nm).

**Chromatin immunoprecipitation**

Chromatin immunoprecipitation (ChIP) was performed using the EZ-Magna ChIP G kit (Merck Millipore) following the manufacturer's instructions but with modifications. Briefly, HUVECs were seeded and transfected in 15 cm dishes. 48 h later, cells were washed twice with PBS, scraped off and fixed with formaldehyde (1% final concentration) for 5 min at RT. After three washing steps, nuclear lysates were prepared as described in the kit's instructions. Nuclei were resuspended in 0.2 mL nuclear lysis buffer and DNA was sheared to a length of ~200 – 700 nt using the Bioruptor Plus sonication device (Diagenode) (10 cycles in high setting 30 sec on/30 sec off, followed by 10 cycles in low setting 30 sec on/30 sec off). ChIP was performed as described in the manufacturer's manual using the following antibodies: anti-MORF4L1/2 (Santa Cruz) (5 µg), and normal mouse IgG (1 µg; supplied with the kit). DNA was recovered using phenol/chlorophorm/isomylalcohol (1:1:1) (Roth) and eluted in 28 µL DNase free water. Results were obtained by qPCR using primers directed against the *FOXO1* promoter region. The sequences can be found in **Supp. Table 5.**

**CRISPR/Cas9**

A dual guide RNA (gRNA) system approach consisting of gRNA-A and gRNA-B was used to remove the triplex-forming region (TFR) of *MIRIAL*. gRNA-A targeted a region upstream of the *Alu* elements harboring the TFR and gRNA-B targeted a region downstream of them. The design of the gRNA was based on the web-interface of CRISPOR (http://crispor.tefor.net/) [3]. The gRNAs were cloned into a lentiCRISPRv2 vector backbone via Esp3I as described in [4]. lentiCRISPRv2 was a gift from Feng Zhang (Addgene plasmid #52961;

http://n2t.net/addgene:52961; RRID:Addgene_52961) [4]. The modification of the lentiCRISPRv2 plasmid with hygromycin resistance was provided by Frank Schnütgen (Dept. of Medicine, Hematology/Oncology, University Hospital Frankfurt, Goethe University, Frankfurt, Germany).

For annealing, the following oligonucleotides were used: *MIRIAL* TFR: gRNA-A: 5′-CAC CGA CCC AGA AGG GAC ACA AGC-3′ and 5′-AAA CGC TTG TGT CCC TTC TGG GTC C-3′; gRNA-B 5′-CAC CGC CCA TCT GAT TTC AAG ATA G-3′ and 5′-AAA CCT ATC TTG AAA TCA GAT GGG C-3‘. gRNA-A was cloned into lentiCRISPRv2 with puromycin resistance, gRNA-B was cloned into lentiCRISPRv2 with hygromycin resistance. After cloning, the gRNA-containing lentiCRISPRv2 vectors were sequenced and purified.

Lentivirus production was performed by transfecting Lenti-X 293T cells (632180, Takara) with GeneJuice Transfection Reagent (70967, Merck), packaging plasmid psPAX2, and the envelope plasmid pVSVG (pMD2.G). psPAX2 and pMD2.G were a gift from Didier Trono (12260, Addgene plasmid, http://n2t.net/addgene:12260; RRID:Addgene_12260 and 12259, http://n2t.net/addgene:12259; RRID:Addgene_12259, respectively). Non-targeting control (NTC) vectors were used as controls.

Viral supernatants were harvested 48 h and 72 h post-transfection, concentrated (Lenti-X Concentrator, Takara) and incubated with HUVECs (p3) in the presence of polybrene (8 µg/mL, Sigma) overnight, followed by expansion and selection (Puromycin 1 µg/mL, Sigma, Hygromycin 100 µg/mL, Merck Millipore). Cells were harvested 10 days post-transduction and total RNA, and genomic DNA were isolated as described elsewhere in this manuscript.

Validation of the CRISPR/Cas9 knockout of the *MIRIAL* TFR was performed from genomic DNA. The CRISPR/Cas9 target site was amplified by PCR using 2x Rapid Taq Master Mix (Vazyme

Biotech) containing forward and reverse primers (10 µM each) and 10 ng purified DNA followed by electrophoresis on an agarose gel stained with Midori Green (Nippon Genetics). The following primers were used: *MIRIAL* TFR KO: 5′- ACT CCT GGC TTC CGG TAT CA -3′ and 5′- AGG CAG GGA GGA TCA GCT ATT -3′; *GAPDH*: 5′- TGG TGT CAG GTT ATG CTG GGC CAG -3′ and 5′- GTG GGA TGG GAG GGT GCT GAA CAC -3′.

**Echocardiography**

Echocardiography was performed to assess cardiac contractile function one, 14 and 28 days after AMI surgery using a Vevo 3100 imaging system (VisualSonics).

**ECIS**

Electrical Cell-substrate Impedance Sensing (ECIS) was performed to assess endothelial barrier integrity and migration after wounding of an endothelial monolayer as described before[19].

**Flow cytometry**

Flow cytometry was used to measure cell cycle progression (Click-iT EdU Flow Cytometry Assay Kit and FxCycle Violet, both Thermo Fisher), DNA damage (anti-H2AX(pS139)-PE, BD Biosciences), mitochondrial mass (Mitotracker Green, Thermo Fisher), mitochondrial membrane potential (TMRM, Thermo Fisher) and reactive oxygen species (ROS) production (MitoSOX, invitrogen) following manufacturer's instructions. Briefly, cells were transfected as described earlier, incubated with the respective labelling compound or antibody for 10-60 min, washed, fixed and permeabilized if required, detached using 0.5% trypsin/EDTA (Gibco), transferred to FACS tubes (BD Biosciences) and then mean fluorescence was measured using the FACS Canto II flow cytometer (BD Biosciences). For viability exclusion cells were stained using either DAPI (Thermo Fisher) or the LIVE/DEAD Fixable Far Red Dead Cell Stain Kit (Thermo Fisher) according to manufacturer's instructions. Unlabeled cells were used as negative control. λ excitation/emission: DAPI 405/450 nm, EdU 488/530 nm, FxCycle Violet 405/450 nm, γH2A.X-PE 488/576 nm, LIVE/DEAD 633/660 nm, MitoSOX 510/580 nm, MitoTracker Green 490/516 nm, TMRM 488/573 nm.

**Genomic DNA isolation**

Isolation of genomic DNA was performed using the PureLink Genomic DNA Mini Kit by Invitrogen according to the manufacturer's instructions. Purified genomic DNA was eluted in a volume of 100 µL of Genomic Elution Buffer (provided with the kit). Concentration of purified genomic DNA was determined using the nanodrop2000 spectrophotometer (Thermo Scientific), diluted to 2.5 ng/µL and then submitted to qRT-PCR. To determine mtDNA content relative to nuclear DNA, we used *MT-ND1* and *RPLP0* primers, respectively. To determine telomere length, we used telomere-specific primers and *36B4D* (single copy gene) primers as reference as described in [5]. All primer sequences can be found in **Supp. Tables 4 and 5**.

**Growth curve**

Cells were seeded in 24-well plates (Greiner), transfected and the cell number was assessed at 0 h, 24 h, 48 h and 72 h post-transfection or only at 0 h and 72 h post-transfection using C-Chip Neubauer Improved disposable hemocytometers (NanoEntek).

**Histology**

Mouse hearts were fixed overnight in 4% PBS-buffered formaldehyde (Histofix, Roth) and subsequently embedded in paraffin. The staining of mouse heart sections with Sirius Red (fibrosis) was carried out as detailed in another study [1]. Infarct size was determined by calculating the ratio of the infarct's length to the circumference of the left ventricle as described before [6]. Fibrosis was assessed by dividing the Sirius red-positive area by the total area of each section. These measurements were performed across three consecutive sections of the hearts.

**Immunoblotting**

Cells were washed with ice-cold PBS (Gibco) and lyzed for 1 h in Radio-Immunoprecipitation Assay (RIPA) buffer (Thermo Scientific) containing 1x protease and phosphatase inhibitors (Halt Protease Inhibitor Cocktail 100×, Thermo Scientific). After centrifugation (10 min, 10,000x *g*), the protein concentration of the supernatant was determined using the Pierce BCA Protein Assay Kit (Thermo Scientific) and equal amounts of protein were boiled in 4x Laemmli Sample Buffer (Bio-Rad) containing β-mercaptoethanol (AppliChem). Samples were separated by SDS-PAGE using Mini-PROTEAN TGX Gels 4-15% (Biorad) and 1x Laemmli Buffer for SDS-PAGE (Serva) and then transferred to Nitrocellulose membrane (Invitrogen) by wet-tank blotting. Membranes were blocked with either 5% non-fat dry powdered milk (Roth) or 5% BSA (AppliChem) in 1x TBS + 0.1% Tween20 (both from Roth). Primary antibodies (Cell Signaling or Novus Bio) are incubated in blocking buffer overnight at 4°C. Secondary antibodies (Agilent Dako) were incubated in TBS-T for 1 h at room temperature. Details on antibodies are found in **Supp. Table 1.** Signal was detected using the Pierce ECL Western Blotting Substrate by Thermo Scientific and the ChemiDoc Touch Imaging System (Biorad). Intensities were measured using Image Lab Software (Biorad) and normalized to the housekeeping proteins GAPDH, β-Tubulin or Vinculin.

**Immunohistochemistry**

Sections for immunohistochemistry were prepared as described before [7] including the antigen retrieval step. Capillary density was visualized using the following primary and secondary antibodies: Isolectin B4 (Vector) and Streptavidin Alexa488 (Invitrogen), wheat germ agglutinin Alexa647 (Thermo Fisher), anti-actin alpha smooth muscle Cy3 (Sigma) and Hoechst 33342 (Anaspec). Slides were mounted using Fluoromount-G mounting medium (Thermo Fisher). Sections were imaged using the Zeiss LSM 780 confocal microscope equipped with a 40x objective and analyzed using Volocity software 6.5.1 (Quorum Technologies).

**KEGG pathway analysis**

The web tool DAVID 6.8 (https://david.ncifcrf.gov/) [8, 9] was used for Kyoto Encyclopedia of Genes and Genomes (KEGG) pathway analysis using the parameters fold change ≥ 1.5 and $p \leq 0.05$.

**miRNA expression levels**

microRNA expression levels were determined using TaqMan assay system by Applied Biosystems. Briefly, samples were reversely transcribed individually for each target miRNA, including U6 snoRNA as a reference using the TaqMan MicroRNA Reverse Transcription Kit. cDNA was then subjected to qRT-PCR using TaqMan MicroRNA Assay probes and TaqMan Fast Advanced Master Mix.

**Mitochondrial structure analysis**

Assessment of mitochondrial fragmentation was performed on confluent HUVECs seeded on 8-well format Ibidi µ-slides following transfection with gapmeR control or gapmeR targeting the lncRNA *MIRIAL*. The cells were subsequently washed and fixed with 4% paraformaldehyde for 10 min at room temperature. Subsequently, HUVECs were permeabilized with 0.2% triton X-100, followed by blocking with 1% human serum albumin (HSA). Mitochondria were stained using rabbit monoclonal TOMM20 antibody 1:100 (Santa Cruz, SC-11415) in 1% HSA for 1 h at 37°C followed by an incubation with FITC-conjugated secondary goat anti-rabbit IgG antibody 1:100 (Thermo Fisher, Alexa Fluor 488, A32731) for 1 h at 37°C. Cells were washed three times with 0.1% HSA and four drops of mounting medium were added into the well. The cells were imaged on a Nikon Eclipse T*i*2-E inverted fluorescence microscope (Nikon Instruments Inc., USA) with a 40X objective. Mitochondrial structure was quantified as described before [10-13]. Briefly, images were opened in FIJI and were adjusted to optimize visualization of the mitochondria (five visual fields were acquired and at least 100 cells were analyzed per condition per experiment; *n*=3 independent experiments). The images were then converted to a binary image and the polygon selection tool was used to outline cells. After selecting the cell, the Analyze Particles function (pixel size 10-infinity; circularity 0.00-1.00) was used to count the number of noncontiguous discrete particles. For every condition, ~100 cells per experiment were analyzed to calculate the mitochondrial fragmentation count (MFC) and form factor (FF) values. MFC is assigned by dividing the number of mitochondrial segments in the cell by the total mitochondrial mass and multiplied by 1000. FF is calculated as (perimeter$^2$/4.π.area).

**Prediction of triplex formation by *MIRIAL***

Potential *RNA*·DNA:DNA triplex formation by *MIRIAL* was predicted using *TriplexAligner* (v1.0) [14]. Briefly, the transcript sequence of *MIRIAL* was used as the RNA input to the algorithm, with the DNA input being made up of promoter or regulatory element sequences. Promoter sequences were defined as being 3500 base pairs upstream and 500 base pairs downstream of transcription start sites as annotated in the R package *TxDb.Hsapiens.UCSC.hg38.knownGene*. The regulatory elements used were taken from *EpiRegio* [15, 16]. *TriplexAligner* predicts triplex formation between RNA and DNA sequences using local alignment with probabilistic scoring matrices learned from triplex-sequencing data. The predicted triplex-forming region of *MIRIAL* was determined by summing the *TriplexAligner* $-\log_{10}(E)$ values obtained across every predicted interaction at base-pair resolution along the length of the transcript. Predicted target sites of *MIRIAL* were further stratified by H3K27ac ChIP-sequencing signal from *ENCODE* [17] (accession ENCFF343KDE, available at s3://encode-public/2020/09/30/aa0c90aa-3f77-4806-8cb1-d98c71c411f6/ENCFF343KDE.bigWig). H3K27ac signal was summed per genomic feature (promoter or REM) and used to identify candidate *MIRIAL* triplex target sites of regulatory importance. Total H3K27ac signal in the feature was multiplied by the *TriplexAligner* $-\log_{10}(E)$ value for the predicted interaction between *MIRIAL* and the respective DNA region to produce a combined score. The logarithm of this score was then rescaled to between 0 and 1 to produce a combined score used to stratify the genomic features.

**RNA affinity purification followed by mass spectrometry**

Nuclear fractions from untransfected HUVECs were prepared using cytoplasmic lysis buffer (10 mM Tris/HCl pH 7.5, 150 mM NaCl, 0.15% (v/v) NP-40), sucrose buffer (10 mM Tris/HCl pH 7.5, 150 mM NaCl, 24% (w/v) sucrose), glycerol buffer (20 mM Tris/HCl pH 7.9, 75 mM NaCl, 0.5 mM EDTA, 0.85 mM DTT, 50% (v/v) glycerol) and nuclei lysis buffer (10 mM HEPES pH 7.6, 7.5 mM $MgCl_2$, 0.2 mM EDTA, 0.3 M NaCl, 1 M urea, 1% (v/v) NP-40, 1mM DTT). The nuclear lysates were adjusted to 150 mM NaCl, 160 U RNase inhibitor, 1 mM DTT, and 1x Protease inhibitor in 1x RNase H buffer (New England Biolabs). For RNA pulldown 200 pmol of biotinylated ASOs directed against *MIRIAL* were incubated overnight at 4°C with pre-cleared (2 h, 4°C) nuclear lysates. The next day, ASO-protein-complexes were captured from the lysate by adding pre-blocked (yeast tRNA and glycogen, 0.2 mg/mL each) and washed (Washing buffer: 50 mM Tris/HCl; pH 8.0, 150 mM NaCl, 1 mM EDTA, 0.05% (v/v) NP-40 and 1x protease inhibitor) magnetic Streptavidin beads (DynaBeads MyOne StreptAvidin C1, Invitrogen) and incubating them for 1 h at 37°C. The beads were then washed (mild washing buffer: 20 mM Tris/HCl; pH 8.0, 10 mM NaCl, 1 mM EDTA, 0.05% (v/v) NP-40 and 1x Protease inhibitor) and eluted using 10 mM d-Biotin in mass spec buffer (10 mM Tris/HCl; pH 8.0, 50 mM NaCl) for 30 min at 37°C and 500 rpm. Parts of the elution fraction were held back for RNA extraction and silver staining. The rest was snap-frozen in liquid nitrogen and then submitted to mass spectrometry.

Eluates from the RNA affinity purification experiments were supplemented with 2 M GdmCl, 50 mM Tris/HCl pH 8.5, 10 mM TCEP and incubated at 95°C for 5 min. Reduced thiols were alkylated with 40 mM chloroacetamid and samples were diluted with 25 mM Tris/HCl pH 8.5, 10% acetonitrile to obtain a final GdmCl concentration of 0.6 M. Proteins were digested with 1 µg trypsin (sequencing grade, Promega) overnight at 37°C under gentle agitation. Digestion was stopped by adding trifluoroacetic acid to a final concentration of 0.5%. Peptides were loaded on multi-stop-and-go tip (StageTip) containing six C18-disks. Purification and elution of peptides was performed as described in Rappsilber and Mann (2007). Peptides were eluted in wells of microtiter plates, dried and resolved in 1% acetonitrile, 0.1% formic acid. Liquid chromatography / mass spectrometry (LC/MS) was performed on Thermo Scientific™ Q Exactive Plus equipped with an ultra-high performance liquid chromatography unit (Thermo Scientific Dionex Ultimate 3000) and a Nanospray Flex Ion-Source (Thermo Scientific).

Peptides were loaded on a C18 reversed-phase pre-column (Thermo Scientific) followed by separation on a with 2.4 µm Reprosil C18 resin (Dr. Maisch GmbH) in-house packed picotip emitter tip (diameter 100 µm, 15 cm from New Objectives) using a gradient from 4% acetonitrile, 0.1% formic acid to 40% eluent B (99% acetonitrile, 0.1% formic acid) for 30 min and a second gradient to 60% B for 5 min with a flow rate 300 nL/min. MS data were recorded by data dependent acquisition. The full MS scan range was 300 to 2000 m/z with resolution of 70,000, and an automatic gain control (AGC) value of 3*E6 total ion counts with a maximal ion injection time of 160 ms. Only higher charged ions (2+) were selected for MS/MS scans with a resolution of 17,500, an isolation window of 2 m/z and an automatic gain control value set to 1*E5 ions with a maximal ion injection time of 150 ms. MS1-Data were acquired in profile mode. MS Data were analyzed by MaxQuant (v1.6.1.0) [19029910] using default settings. Proteins were identified using reviewed human reference proteome database UniProtKB with 71785 entries, released in 2/2018. The enzyme specificity was set to Trypsin. Acetylation (+42.01) at N-terminus and oxidation of methionine (+15.99) were selected as variable modifications and carbamidomethylation (+57.02) as fixed modification on cysteines. False discovery rate (FDR) for the identification protein and peptides was 1%. Label free quantification (LFQ) and intensity-based absolute quantification (IBAQ) values were recorded. Data were further analyzed by Perseus (v. 1.6.1.3). Contaminants and reversed identification were removed. Protein identification with at least 4 valid quantification values in at least one group were further analyzed. Missing values were replaced by background values from normal distribution. Student´s *t* test was used to identify significantly enriched proteins between experimental groups.

**RNA immunoprecipitation**

For RNA immunoprecipitation, 5 µg of antibodies (p53 (1C12) (Cell Signaling) or MORF4L1/2 (Santa Cruz) and Normal Mouse IgG (Santa Cruz)) were used. Briefly, HUVECs cultured on 150mm-dishes (Greiner) until confluency, crosslinked with 50 mJ/cm$^2$ UV light, lyzed in lysis buffer (50 mM Tris/HCl pH 8.0, 150 mM NaCl, 0.5% NP40, 1 mM EDTA, protease inhibitors) and incubated overnight with washed antibody-coated magnetic protein G beads (Dynabeads, invitrogen) in lysis buffer without NP40. Beads were then washed with binding buffer (50 mM Tris/HCl pH 8.0, 150 mM NaCl, 0.05% NP40, 1mM EDTA), eluted by proteinase K digest (New England Biolabs) and RNA was recovered using phenol/chloroform/isoamylalcohol (1:1:1) (Roth). 5% of the input and the elution fractions were held back, boiled in 4x Laemmli Sample Buffer (Bio-Rad) containing β-mercaptoethanol (AppliChem) and used for subsequent SDS-PAGE and immunoblotting.

**RNA interference & miRNA overexpression**

HUVECs were cultured at 50-60% confluency before being transfected with siRNA (QIAGEN or Sigma), LNA GapmeRs (QIAGEN) or pre-miRs (ambion) according to manufacturer's recommendations using Lipofectamine RNAiMax (Invitrogen) and serum reduced OptiMEM (Gibco). LNA control A (QIAGEN), siRNA directed against Firefly Luciferase (siCtrl) (Sigma) or pre-miR negative control (ambion) were transfected as controls. Details on antisense oligonucleotides (ASOs) and miRNA precursors can be found in **Supp. Tables 2 and 3**, respectively. 4 h after transfection the medium was changed to full endothelial growth medium.

**Seahorse**

48 h prior to the assay, HUVECS were transfected with either LNA Control or LNA *MIRIAL*. 24 h after the transfection, cells were re-seeded into Seahorse XF96 Cell Culture Microplates

(Agilent) coated with fibronectin and gelatin. The Seahorse XF Glycolysis Stress Test and Seahorse XF Cell Mito Stress Test were performed using the same-named kits and the Seahorse XFe96 Analyzer (all by Agilent) following the manufacturer's instructions. Cells were assayed in Seahorse XF Base Medium (Agilent) containing L-glutamine, D-glucose, and sodium pyruvate (all from Sigma), pH 7.4. Oxygen consumption rate (OCR) and extracellular acidification rate (ECAR) were normalized to cell number per well determined by Hoechst 33258 staining after the assay. Fluorescence was measured using the Glomax Multi+ Detection System plate reader by Promega (fluorescence: excitation 365 nm, emission 410-460 nm; absorption: 560 nm).

**Senescence-associated β-galactosidase staining**

Cellular senescence was assessed using the Senescence Associated β-Galactosidase Staining kit (Cell Signaling) 48 hours post-transfection. Cells were washed twice with PBS (Gibco) and fixed for 5 minutes at room temperature using 1x fixative solution (20% formaldehyde, 2% glutaraldehyde in PBS). After three washes with PBS (Gibco), cells were incubated with staining solution containing the chromogenic substrate 5-bromo-4-chloro-3-indolyl-β-D-galactopyranoside (X-Gal) (pH adjusted to pH 6.0 with HCl) overnight at 37°C without $CO_2$ in a non-humidified incubator. Images were captured at 5x magnification using a Zeiss Axiovert 100.

**Sequence homology alignment**

Sequence alignment of human and mouse *MIRIAL/Mirial* was performed as described in [18]. The LALIGN DNA:DNA tool of the FASTA Sequence Comparison software (University of Virginia) was used with default parameters [19]**.**

**Spheroid sprouting assay**

HUVECs were transfected with either LNA Control or LNA *MIRIAL*. The next day, 400 cells per well are re-seeded in semi-solid medium (80% EBM + 20% methylcellulose (0.6 g/L, Sigma)) in 96-well U-bottom plates (Greiner) to form three-dimensional spheroids. After additional 24 h, the spheroids were carefully harvested, centrifuged, re-suspended in a mixture of 80% methocoel and 20% FBS (Gibco) and then re-seeded in a collagen medium (10x M199 (Sigma), collagen stock solution type I (Corning), HEPES, NaOH (both Sigma). After a 30 min incubation at 37°C, the embedded spheroids are stimulated with VEGF-A (PeproTech) at a final concentration of 50 ng/mL in endothelial growth medium. Unstimulated spheroids fed with only endothelial growth medium were included as controls. 24 h later, spheroids were fixed with HistoFix (Roth) for 30 min and imaged using the Zeiss AxioVert microscope with 10x magnification. Cumulative sprout length was quantified using Image J software.

**Subcellular fractionation**

Cytoplasmic, nucleoplasmic and chromatin fractions of untransfected HUVECs were isolated as described in[19].

**Total RNA isolation, cDNA synthesis and qRT-PCR**

Total RNA was isolated using QIAzol lysis reagent (QIAGEN) and the Direct-zol RNA MiniPrep Kit (Zymo Research) (for HUVEC samples) including recommended DNase-I digestion, or the miRNeasy Micro Kit or Mini Kits (QIAGEN) (for in vivo samples) following the manufacturer's instructions. RNA was reversely transcribed to cDNA using random hexamer primers (QIAGEN) and MultiScribe Reverse Transcriptase (Invitrogen). qRT-PCR was performed using Fast SYBR Green Master Mix (Applied Biosystems) and either Step One Plus or ViiA 7 thermocyclers (Applied Biosystems). Human and mouse primer sequences can be found in **Supp. Tables 4 and 5**, respectively. Gene expression was normalized to Ribosomal protein lateral stalk subunit P0 (*RPLP0*) or glyceraldehyde-3-phosphate dehydrogenase (*GAPDH*) expression. Gene expression levels were analyzed using the $2^{-\Delta CT}$ method.

### 3. Supplementary Tables:

**Supp. Table 1: Antibodies**

| Target antigen | Supplier | Catalogue # | Working concentration |
|---|---|---|---|
| Actin α, smooth muscle, Cy3 | Sigma-Aldrich | C6198 | 1:300 |
| AMPKa (D63G4) | Cell Signaling | 5832 | 1:1000 |
| c-Myc | Cell Signaling | 9402 | 1:500 |
| FOXO1 (C29H4) | Cell Signaling | 2880 | 1:1000 |
| GAPDH (14C10) | Cell Signaling | 2118 | 1:2000 |
| Goat anti-mouse HRP | Agilent Dako | P0447 | 1:2000 |
| Goat anti-rabbit Alexa Fluor 488 | Thermo Fisher | A32731 | 1:100 |
| Goat anti-rabbit HRP | Agilent Dako | P0448 | 1:2000 |
| H2AX (pS139) PE | BD Bioscience | 562377 | 1:100 |
| Mitochondrial Dynamics Antibody Sampler Kit II | Cell Signaling | 74792 | 1:1000 |
| MORF4L1/2 | Santa Cruz | sc-514659 | 5 µg |
| MORF4L2 | Novus Bio | NBP2-47370 | 1:1000 |
| Normal mouse IgG | Santa Cruz | sc-2025 | 5 µg |
| p21, WAF1, CIP1 (12D1) | Cell Signaling | 2947 | 1:1000 |
| p53 | Cell Signaling | 9282 | 1:500 |
| p53 (1C12) | Cell Signaling | 2524 | 5 µg |
| Phospho-AMPKa (Thr172) (D4D6D) | Cell Signaling | 50081 | 1:1000 |
| phospho-FOXO1 (Ser256) (E1F7T) | Cell Signaling | 84192 | 1:1000 |
| Phospho-p53 (Ser15) | Cell Signaling | 9284 | 1:1000 |
| TAF15 | Cell Signaling | 28409 | 1:1000 |
| TOM20 | Santa Cruz | sc-11415 | 1:100 |
| Vinculin XP (E1E9V) | Cell Signaling | 13901 | 1:2000 |
| β-Tubulin (9F3) | Cell Signaling | 2128 | 1:2000 |

**Supp. Table 2: Antisense oligonucleotides**

| ASO | Concentration | Sequence |
|---|---|---|
| LNA *MIRIAL* | 20 nM | GGCATTTAACGGTCTG |
| siRNA Control | 10-60 nM | CGUACGCGGAAUACUUCGA |
| siRNA MORF4L2 | 10 nM | AGGGTTGAATGAGTTCCAGAA |
| siRNA *MYC* | 60 nM | GATCCCGGAGTTGGAAAACAA |
| siRNA *p53* | 10 nM | CAGCATCTTATCCGAGTGGAA |
| siRNA *TAF15* | 10 nM | TACGGTGGAGACCGAAGTGGA |

**Supp. Table 3: miRNA precursors for miRNA overexpression (pre-miRs by ambion)**

| miRNA precursor | Concentration | Mature sequence |
|---|---|---|
| Hsa-miR-23a-3p | 10 nM | AUCACAUUGCCAGGGAUUUCC |
| Hsa-miR-27a-3p | 10 nM | UUCACAGUGGCUAAGUUCCGC |
| Hsa-miR-24-3p | 10 nM | UGGCUCAGUUCAGCAGGAACAG |

**Supp. Table 4: Human primers for q(RT)-PCR**

| Target | Strand | Sequence |
|---|---|---|
| *36B4D* | Fwd | CCCATTCTATCATCAACGGGTAC |
| *36B4D* | Rev | CAGCAAGTGGGAAGGTGTAATCC |
| *CDKN1A* | Fwd | GCGACTGTGATGCGCTAATG |
| *CDKN1A* | Rev | GAAGGTAGAGCTTGGGCAGG |
| *CMBL* | Fwd | CAGACTTCTTTGTAGGGCAAGAG |
| *CMBL* | Rev | GGCATGACACTGTTGTTTCAGA |
| *CYFIP2* | Fwd | TTCCGTATCCACCGTCCAAT |
| *CYFIP2* | Rev | CGCTGGGTAATGAGTCTGTTC |
| *FOXO1* | Fwd | GTCAAGACAACGACACATAG |
| *FOXO1* | Rev | AAACTAAAAGGGAGTTGGTG |
| *FOXO1* promoter | Fwd | GAGATTTGGGGGAACGAAGC |
| *FOXO1* promoter | Rev | CGAGGAGCCTCGATGTGGAT |
| *GADPH* | Fwd | GAGTCAACGGATTTGGTCGT |
| *GADPH* | Rev | TTGATTTTGGAGGGATCTCG |
| *GDF15* | Fwd | TAACCAGGCTGCGGGCCAAC |
| *GDF15* | Rev | CAGCCGCACTTCTGGCGTGA |
| *HMOX1* | Fwd | AGTCTTCGCCCCTGTCTACT |
| *HMOX1* | Rev | CTTCACATAGCGCTGCATGG |
| *LIF* | Fwd | TACGCCACCCATGTCACAAC |
| *LIF* | Rev | CTTGTCCAGGTTGTTGGGGA |
| *MDM2* | Fwd | CGAGCTTGGCTGCTTCTGG |
| *MDM2* | Rev | GTACGCACTAATCCGGGGAG |
| *MIRIAL* | Fwd | CCTAGCAGCCAGTTACCCAA |
| *MIRIAL* | Rev | TCCCGGCAAGTTAGAAAGCT |
| *MIRIAL* TFR KO | Fwd | ACTCCTGGCTTCCGGTATCA |
| *MIRIAL* TFR KO | Rev | AGGCAGGGAGGATCAGCTATT |
| *MORF4L2* | Fwd | AGAAGGAAGATTGTTGGTTG |
| *MORF4L2* | Rev | GCTGGTTCAAGATTTTTCTG |
| *MT-ND1* | Fwd | CCCTAAAACCCGCCACATCT |
| *MT-ND1* | Rev | GAGCGATGGTGAGAGCTAAGGT |
| *MYC* | Fwd | AGCTGCTTAGACGCTGGATTTT |
| *MYC* | Rev | TCGAGGTCATAGTTCCTGTTGG |
| *PRKAA1* | Fwd | TGTGATGGGATCTTCTATACC |
| *PRKAA1* | Rev | CCCTGATATCTTTGATTGTGG |
| *RPLP0* | Fwd | TCGACAATGGCAGCATCTAC |

| | | |
|---|---|---|
| *RPLP0* | Rev | ATCCGTCTCCACAGACAAGG |
| *RRM2B* | Fwd | AGTTCTCGCCGGTTTGTCAT |
| *RRM2B* | Rev | AAGGAAGCCTGTGCCTGTTT |
| *TAF15* | Fwd | CAAAGCTATTCTGGCTATGG |
| *TAF15* | Rev | CTGGTTATTATATGGTTGCTGG |
| *TBP* | Fwd | GAGCTGTGATGTGAAGTTTCC |
| *TBP* | Rev | TCTGGGTTTGATCATTCTGTAG |
| Telomere | Fwd | GGTTTTTGAGGGTGAGGGTGAGGGTGAGGGTGAGGGT |
| Telomere | Rev | TCCCGACTATCCCTATCCCTATCCCTATCCCTATCCCTA |
| *TP53* | Fwd | TTTCACCCTTCAGATCCGTGG |
| *TP53* | Rev | AGTCTGAGTCAGGCCCTTCT |
| *TP53I3* | Fwd | TGGCTCTTGATGGTCGATGG |
| *TP53I3* | Rev | AGGCAGAATTTGCTCCGTGA |

**Supp. Table 5: Mouse primers for q(RT)-PCR**

| Target | Strand | Sequence |
|---|---|---|
| *Rplp0* | Fwd | GCGTCCTGGCATTGTCTGT |
| *Rplp0* | Rev | GAAGGCCTTGACCTTTTCAGTAAG |
| *Mirial* KO | Fwd | GCTTTAGGCCCTCTAACGGT |
| *Mirial* KO | Rev | CTCGCTGTCACCAGCAGAAA |
| *Mt-Nd1* | Fwd | CTAGCAGAAACAAACCGGGC |
| *Mt-Nd1* | Rev | CCGGCTGCGTATTCTACGTTA |
